# CIViC-Fact: a proof-of-concept framework for AI-assisted verification of cancer variant interpretations

**DOI:** 10.1101/2025.09.10.675443

**Authors:** Caralyn Reisle, Cameron J. Grisdale, Kilannin Krysiak, Arpad M. Danos, Mariam Khanfar, Erin Pleasance, Jason Saliba, Melika Hanos, Nilan V. Patel, Asmita Jain, Morteza Seifi, Joshua F. McMichael, Ajay C. Venigalla, Malachi Griffith, Obi L. Griffith, Steven J. M. Jones

**Author notes:** corresponding authors, address correspondence to: Obi L. Griffith,; Steven J. M. Jones.

## Abstract

Accurate interpretation of genomic variants is critical for precision oncology but remains slow and dependent on specialized expertise. Public knowledgebases such as the Clinical Interpretation of Variants in Cancer (CIViC) help by curating literature-backed variant interpretations in a structured form, yet verification and review have become major bottlenecks. Large language models (LLMs) offer a potential mechanism for accelerating biomedical claim verification, but their rapid turnover, variable availability, and known risks of unsupported reasoning require standardized and reproducible evaluation before integration into curation workflows. To address this, we developed CIViC-Fact, an expert-curated, full-text benchmark and evaluation framework. Domain experts linked structured cancer-variant claims to sentence-level evidence from source publications, including evidence from full-text articles, tables, and non-abstract sections that are commonly omitted from existing biomedical question-answering and scientific fact-checking datasets. Claim-verification reference labels were derived from CIViC records, revision histories and controlled data augmentation. A major finding of CIViC-Fact is that abstracts are insufficient for realistic biomedical claim verification. In the evaluated development subset of text-verifiable entries with full-text access, fewer than 30% could be fully validated from the abstract alone, highlighting the importance of full-text evaluation for biomedical curation.

Upon the application of our fact-checking pipeline to newly submitted CIViC entries, after excluding entries requiring supplementary material or images for validation, automated retrieval successfully identified appropriate evidence for most cases (93%), supporting low-incremental-effort evaluation of future systems. Fine-tuning improved agreement with CIViC-Fact reference labels on the static benchmark, but larger general-purpose models performed better on a heterogeneous post-cutoff cohort. These findings support CIViC-Fact primarily as a reproducible framework for comparing evolving retrieval and verification systems rather than as validation of a single deployment-ready model. These findings suggest that, in a rapidly changing model landscape, the durable contribution is not a single optimized model but a reproducible benchmark framework that enables continual testing, model substitution, and lightweight updating through small high-quality few-shot exemplar sets.

## Introduction

Tailoring treatment to an individual’s genetic profile through precision oncology has played an increasing role in cancer treatment over the last decade. Research has shown value in approaching cancer on a case-by-case, personalized basis^1–5^. However, its translation into standard of care remains constrained. The major obstacle lies in variant interpretation. Analysis requires deep expertise in cancer biology, genomics, and bioinformatics^6^ making it slow, resource-intensive, and difficult to scale

To mitigate repetitive case-by-case literature review, knowledgebases such as OncoKB^7^, CIViC^8,9^, and others^10–17^ curate structured, literature-backed variant annotations, enabling programmatic access to actionable insights^18^. Among these, CIViC (Clinical Interpretation of Variants in Cancer) is unique in being openly accessible, crowdsourced, and allowing anyone to contribute or edit entries enabling rapid growth through community contributions. Yet, ensuring the accuracy and auditability of data that may inform treatment decisions remains paramount^19^. Content must be reviewed by a small team of expert editors, and this verification stage has become the rate-limiting step, creating a new major bottleneck in knowledgebase curation.

Natural language processing (NLP), the AI-driven analysis of human language, has been applied to several precision oncology applications, including variant classification^20^, relation extraction^21–23^, and others^24–26^. However, biomedical curation is a high-stakes, evidence-grounded task: a useful system must not only generate a plausible answer, but connect structured claims to the relevant source evidence and identify when a claim requires revision. Large language models (LLMs) show promise in generalizing to biomedical domains^27^. However, their unreliability (hallucinations, factual errors)^28^, and lack of transparency makes them unsuitable for deployment without rigorous evaluation.

Existing benchmarks do not adequately capture this setting. Most fact-checking datasets focus on generall^29,30^ or political^31^ claims and are unsuitable for fact-checking in a scientific context. Scientific publications are particularly challenging, as they rely on technical language, jargon, and domain-specific expertise that make it difficult for NLP models to verify claims. Existing scientific fact-checking datasets are further limited by relying primarily on abstracts, overlooking claims that can only be verified using full-text evidence^32,33^. This limitation is critical: Cancer variant curation often requires evidence from full-text articles, including results, methods, tables or multiple non-contiguous passages. This creates both an intra-document retrieval problem and a claim-verification problem, compounded by structured biomedical terminology and curation conventions and ontologies^34^ that evolve over time. Reproducible evaluation is therefore needed to determine which retrieval and language-model components are suitable for expert-assisted curation and to reassess them as models and curation practices change.

To address this gap, we present CIViC-Fact, an expert-curated, full-text dataset and proof-of-concept framework for AI-assisted biomedical curation. Domain experts manually linked structured CIViC claims to sentence-level evidence from their source publications, enabling separate evaluation of passage retrieval, evidence-grounded claim verification and candidate revision generation. Unlike scientific fact-checking datasets that rely primarily on abstracts, CIViC-Fact includes evidence from non-abstract sections and tables while preserving the structured CIViC data model. We use this framework to assess where AI can supplement expert curation by surfacing relevant evidence, triaging claims and proposing candidate revisions, while retaining expert review as the final decision point. The resulting dataset also provides reproducible infrastructure for comparing retrieval and verification systems as models and curation standards evolve.

## Methods

### Collection of Sentence Level Evidence Provenance

CIViC collects variant interpretations from literature stored as *Evidence Items*. Each *Evidence Item* in CIViC links a scientific article to a variant interpretation with predictive (therapeutic), diagnostic, prognostic, predisposing, oncogenic or functional significance. CIViC *Evidence Items* include both positive and negative findings depending on the *Evidence Direction* (supports or does not support respectively). CIViC-Fact was constructed by aligning CIViC’s curated *Evidence Items* (Fig 1a), hereafter referred to as claims, with the full-text scientific articles from which they were originally derived.

**Figure 1.**
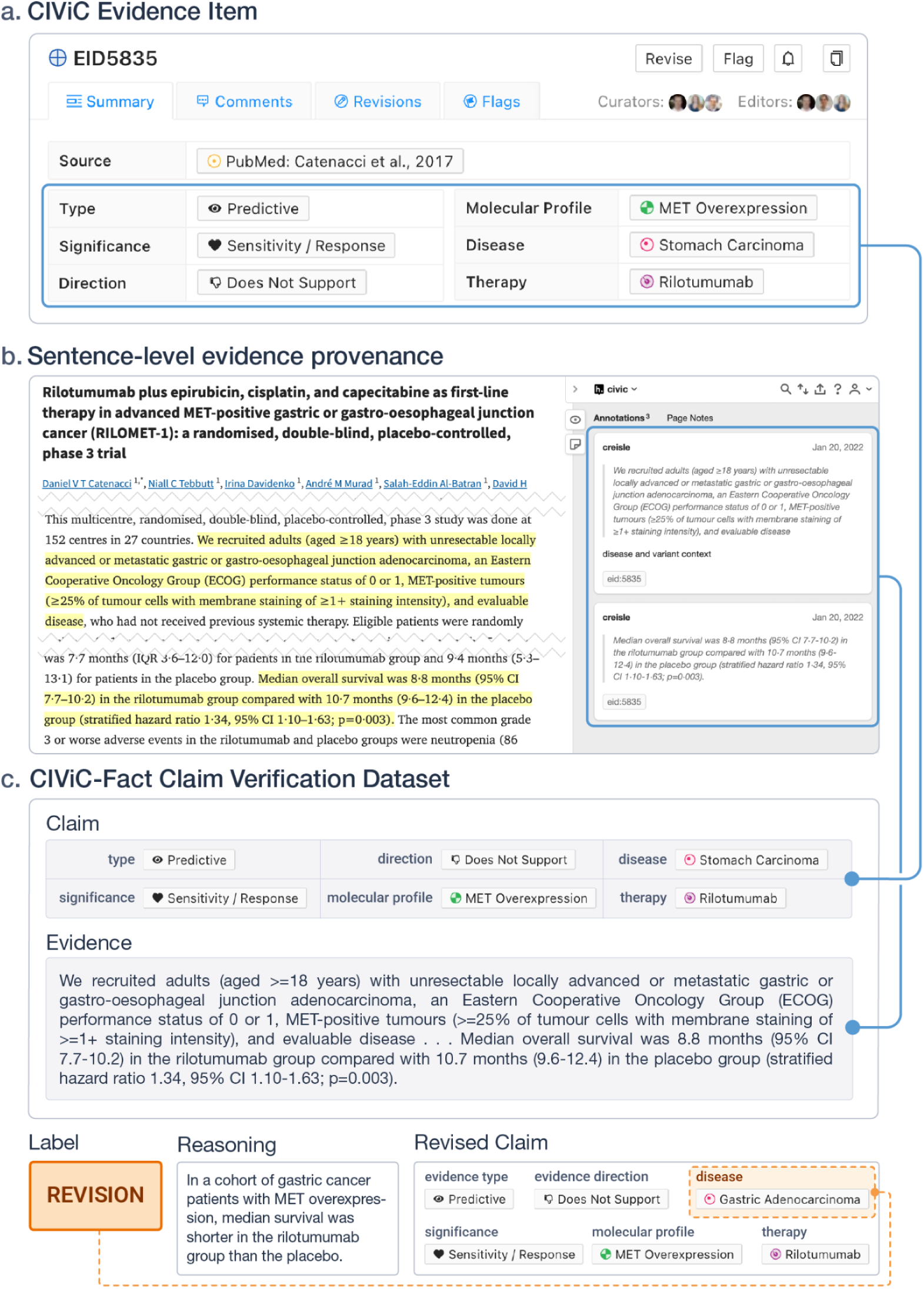
An example entry in the CIViC-Fact dataset. A) The claim is composed from the structured elements of a CIViC entry: molecular profile (gene and variant), disease, therapy, type, significance, and phenotypes. B) Evidence was selected from PubMed Central (PMC5898242) by a curator using the web-tool hypothes.is. C) The claim in the CIViC-Fact dataset is composed of the structured elements of the CIViC entry (except evidence level and rating) and the evidence is manually curated excerpts from the source manuscript. In this example, the label is derived from the revision history in CIViC where the disease field was revised from “stomach carcinoma” to “gastric adenocarcinoma”.

CIViC-Fact augments claims in the CIViC knowledgebase with sentence-level evidence provenance, linking each claim to the specific sentence spans in the article that provide the evidence used to evaluate it using the tool hypothes.is (Fig 1b). Claims that require evidence outside the main article text were tagged as “requires-supplement” or “requires-image” where appropriate. After data was collected by annotators, it was processed and formatted. Tagged annotations were downloaded via the hypothes.is API along with CIViC entries via the CIViC API. Articles were then matched to their full-text XML version from pubmed/PMC based on the article referenced by the CIViC entry. XML articles were converted to the standard Bioc format^35^ using the parsing package developed as part of CIViCmine^26^ (https://GitHub.com/jakelever/biotext). The Bioc format was not used directly from PMC as it excluded elements of interest. Text from the hypothes.is annotations were matched to text in the Bioc articles (removing whitespace errors and other formatting issues from the data collected by hypothes.is).

Claim verification is a crucial step in fact-checking where the verifier is given both the claim and relevant evidence (extracted sentence spans) and asked to label the stance of the evidence toward the claim. Existing stance-classification and fact-checking datasets typically use either two labels (SUPPORTS, DOES NOT SUPPORT) or three labels (SUPPORTS, REFUTES, NOT ENOUGH INFORMATION)^30,32,36–38^. Because CIViC-Fact is intended to flag and potentially correct problematic claims, we introduce a REVISIONS label for cases in which the evidence would support the claim after a limited modification (Supplementary Table 1). For example, consider a source article that evaluates EGFR mutations in relation to erlotinib but finds no evidence of sensitivity. In a traditional three-label framework, the statement “EGFR mutations are associated with erlotinib sensitivity” would be labelled REFUTES. In CIViC-Fact, however, evidenceDirection is part of the structured claim. If the claim specifies evidenceDirection = SUPPORTS, the expected label is REVISIONS, with the proposed correction changing evidenceDirection to DOES NOT SUPPORT. If the claim already specifies evidenceDirection = DOES NOT SUPPORT, the expected label is SUPPORTS because the source evidence is consistent with the complete structured claim.

NEI is assigned only when the available evidence is insufficient to evaluate the claim, is unrelated to the claim, or would require construction of a substantially different claim rather than a revision. It does not indicate that the evidence contradicts or fails to support the biological assertion. When the source directly addresses the claim, including evidence in the opposite biological direction, that relationship is represented by the “evidenceDirection” field in the input claim. Claims requiring supplementary data or interpretation of figures are also assigned NEI because the current verification framework is text-only; future work may extend the task to multimodal evidence as sufficient examples become available for evaluation. Otherwise, claim–evidence pairs annotated in Hypothes.is are assigned SUPPORTS by default, and documented CIViC revision histories are used to construct REVISIONS pairs (Fig. 1c). To broaden the claim-verification task, we additionally generate examples through controlled modification of reference claims^39^ as well as fully-generated examples^38^.

### Reference-label construction and provenance

There are a number of different ways errors can happen in CIViC and other cancer knowledgebases beyond simple curator mistakes. There is a level of nuance where a curator may enter data that is not completely incorrect but would be better represented by a different ontology term or is missing a concomitant variant, etc. In order to train a discriminator model that is sensitive to such changes we augment the CIViC-Fact dataset in several ways.

In order to generate claim/evidence pairs which represent common curator errors in CIViC curation, we generate REVISIONS examples by modifying different fields within the claim: disease, therapies, molecular profile, etc. (Fig 2). CIViC uses the disease ontology^34^ to curate disease terms. Since the disease ontology includes hierarchical disease relationship information, we are able to automatically modify the existing disease term in a claim with a more or less specific term which would require revision if entered in CIViC. Similarly, we use NCIt to find more or less specific drug terms.

**Figure 2.**
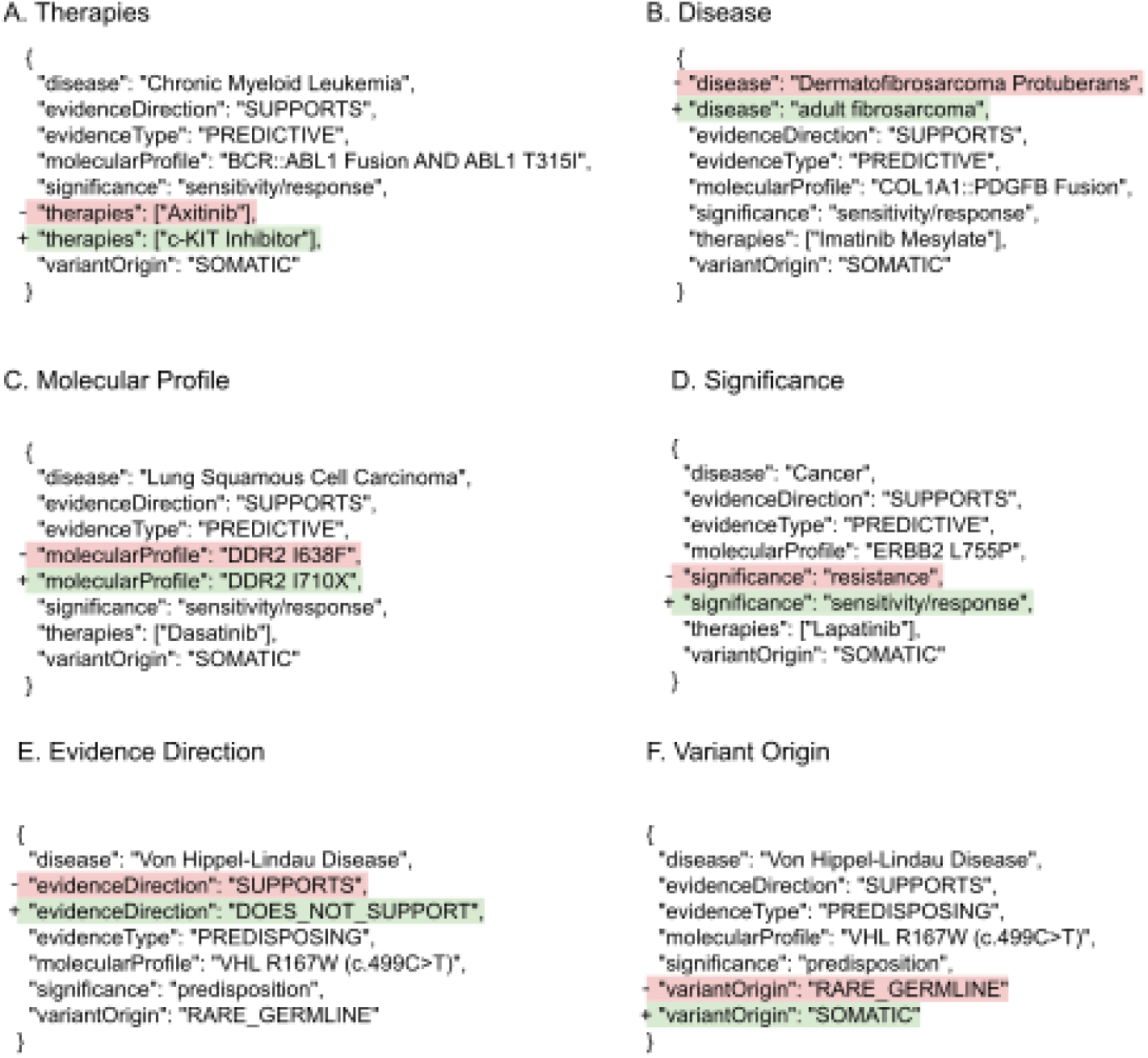
Examples of generated claims requiring revision. The original CIViC claim was modified, when the modified claim is given as input we expect the model to suggest the original claim as the correct revision.

Claims may be supported by different parts of the paper and evidence supporting a claim may be rewritten or summarized in several different ways that are all valid. Curators adding content to CIViC curate a written statement along with the structured elements summarizing the evidence in the paper related to the claim. Naturally these statements do not always mention all elements of the claim as they are partly redundant with the structured elements. We include the structured claim paired with the written statement as additional claim verification pairs. However, we first process these written statements with PubTator^40^ and matching via regular expressions to determine if the written statement is sufficient evidence for the structured claim and we include it as a negative NEI example when it is not.

It is also possible for human curators to enter content incorrectly, misunderstand the CIViC data model, or even link to the wrong source document. Therefore we generated additional negatives to represent such cases by creating plausible claims based on the variants, diseases, and therapies found in the paper (via PubTator^40^) randomly paired with *Evidence Type* and *Significance*. The generated claims were then paired with sentences in the paper using the biomedical passage retrieval model, Specter^41^. The amount of evidence was chosen to match the distribution of selections curated in the expert-selected evidence.

Claim/Evidence Pairs were partitioned into training (∼80%), development (∼10%) and test (∼10%). Pairs were partitioned by stratified random sampling to ensure representation of all document and claim types. Stratification included document journal and *evidence type/significance*. A document was limited to a single partition so claims from the same document would be in the same partition to avoid possible information leakage due to shared evidence selections across claims. Similarly, pair IDs were kept across dataset builds, and manuscripts were kept within a partition to avoid information leakage. In order to avoid bias whereby claim verification models can learn patterns within the claim alone^42^ we generate many negatives and subsample them based on reducing the bigram bias^43^ toward any given label within a partition.

### Passage-retrieval evaluation

To support fact-checking of new CIViC claims, the pipeline requires an automated method for selecting relevant passages from the linked source publication rather than relying on manual evidence identification. Because each CIViC claim is already associated with a specific paper, passage retrieval was formulated as an intra-document ranking task: given a claim and passages from its source article, the model ranks the passages by their relevance to the claim. The fine-grained traceability built into the CIViC-Fact dataset enables systematic evaluation of embedding models^41,44–46^, often used for passage retrieval in retrieval-augmented generation (RAG^47^) pipelines, for their effectiveness in selecting appropriate supporting passages from source articles. The highest-ranked passages are then provided as evidence to the claim-verification model (Fig. 3).

**Figure 3.**
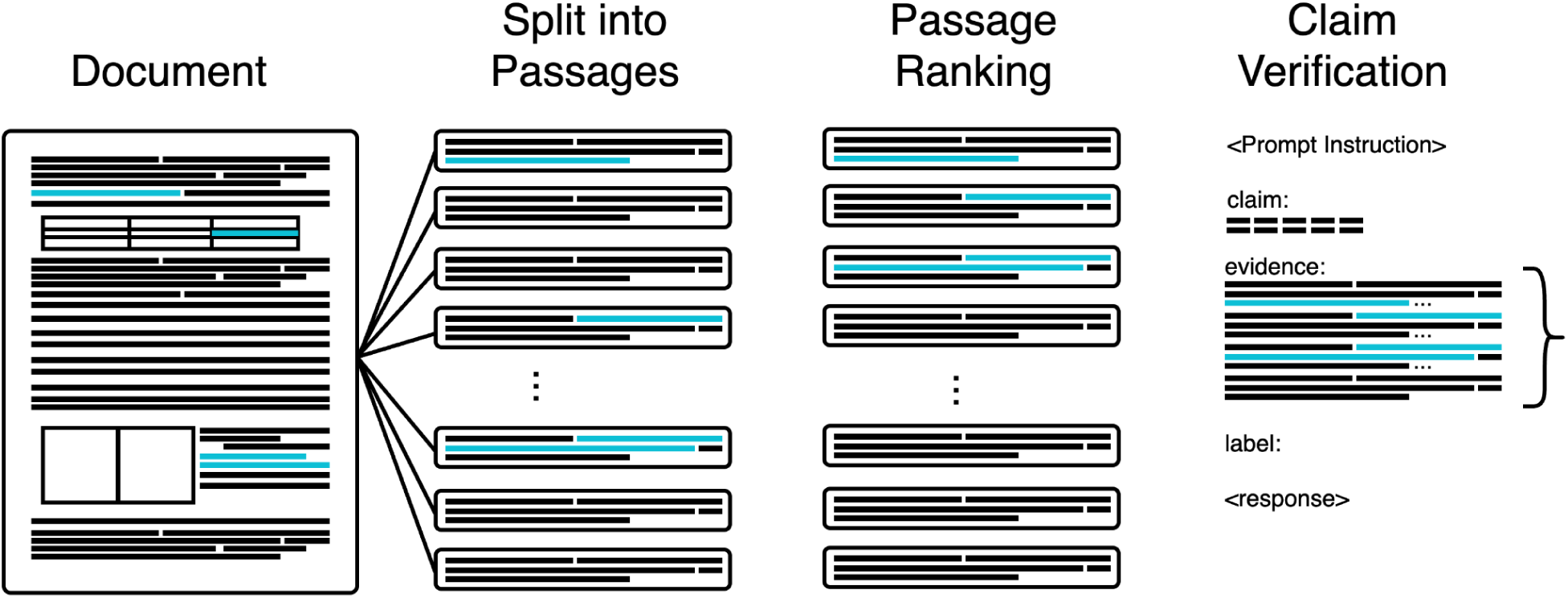
Retrieval of relevant document passages as evidence for claim verification. Relevant selections are shown in blue. A document is split into passages which may or may not overlap content relevant to the claim in question. A passage ranking model is used to order the chunks according to their relevance to the claim. An ideal ranking will put all passages overlapping relevant content at the top of the ranking. The claim and the top-k passages are concatenated as input for claim verification.

CIViC claims comprise multiple structured fields rather than a single natural-language statement. The molecular profile (gene and variant), disease, therapy, significance, variant origin, therapy interaction type, evidence direction and phenotypes were therefore converted into textual representations for retrieval. Five linearization strategies were evaluated: a natural-language template, semicolon-delimited field–value pairs, a tab-delimited two-row table, JSON and a values-only list (Fig. 4)^48^. These comparisons were performed as part of the passage-retrieval evaluation. Claim verification used the JSON representation throughout.

**Figure 4.**
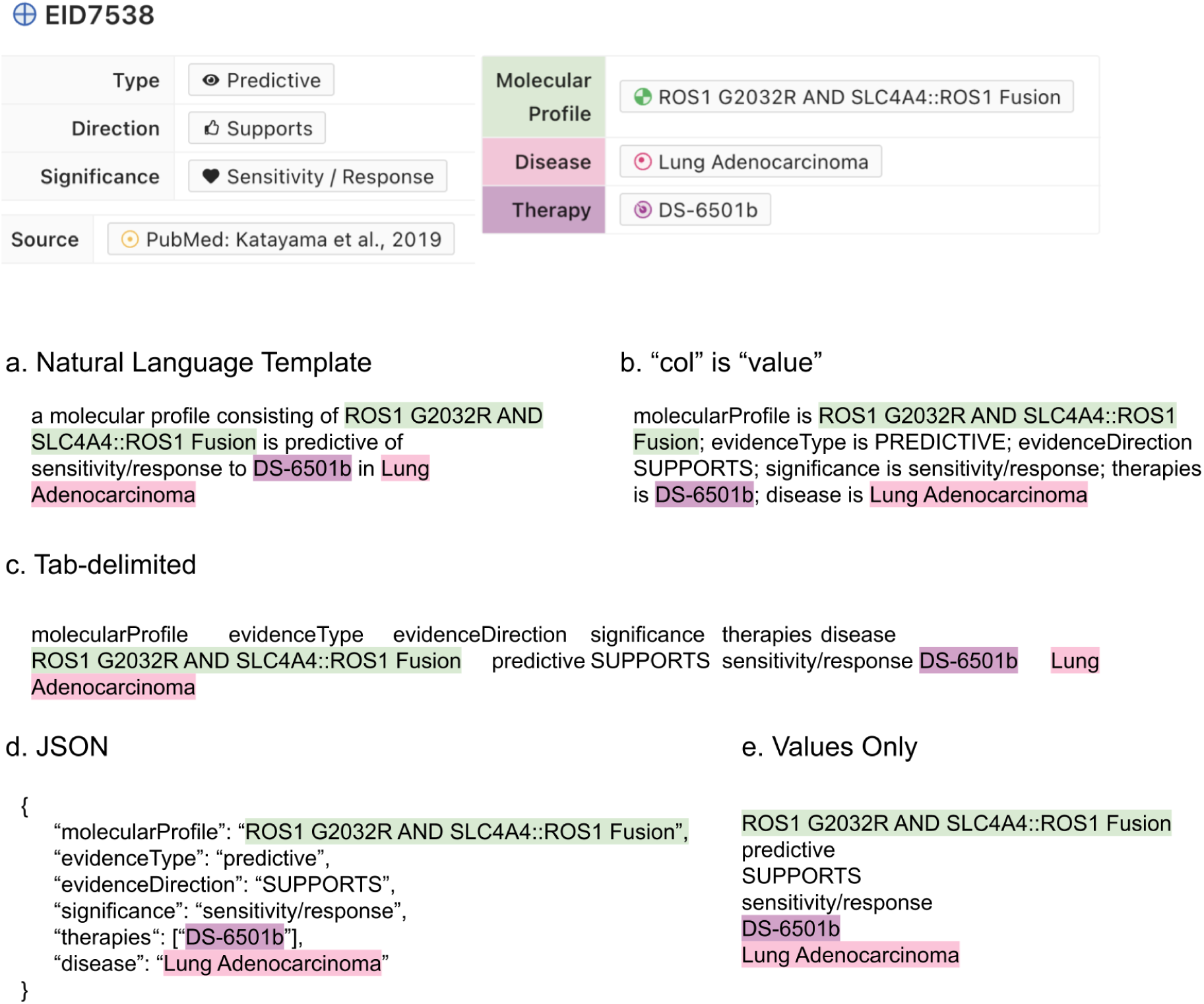
Claim-linearization strategies. Some claim details are omitted for space. (a) Natural-language representation generated using predefined templates. (b) Semicolon-delimited field–value pairs following the FEVEROUS convention. (c) Tab-delimited two-row table containing field names and corresponding values. (d) JSON object containing field–value pairs. (e) Values-only representation, with one value per line.

From the CIViC-Fact dataset, only provable claims (excludes NEI or claims with conflicting evidence) with access to the full text were used in evaluating passage retrieval. Hereafter, we will refer to this as the CIViC-Fact retrieval subset (n_train_=3456, n_dev_=1194, n_test_=1120). We generated passages by preprocessing manuscripts and splitting them into 1000 character passages with a 100 character overlap. Claim/passage pairs were then ranked and the top-k passages were selected as evidence. The resulting prompt context was provided to the LLM along with a prompt instruction for response generation (Fig 3). We evaluated several popular general and biomedical-specific embedding models (Specter2^49^; MedCPT^44^;

Qwen3-Reranker^50^; and BGE reranker^45^) in addition to the BM25^51^ standard baseline. Each retrieval model received a linearized claim and document passages from its linked article and ranked the passages by relevance. Although it is more efficient, bi-encoding (encode the claim and passages separately), degraded performance drastically (data not shown) and it is not necessary as we are only encoding the passages of a single paper at a time, therefore we chose to evaluate cross-encoding models specifically. We evaluated the performance using mean average precision (mAP^52^). mAP measures both the relevance of recommended items and the quality of their ranking.

### Claim-verification evaluation

JSON was chosen as the method for claim linearization for claim verification as it allows the simplest comparison to the potentially revised claim and the least alteration from the initial CIViC format. For evidence construction, the matched text from the hypothes.is annotations were concatenated in order of appearance, with ellipsis delimiters. The general format is given in Figure 5.

**Figure 5.**
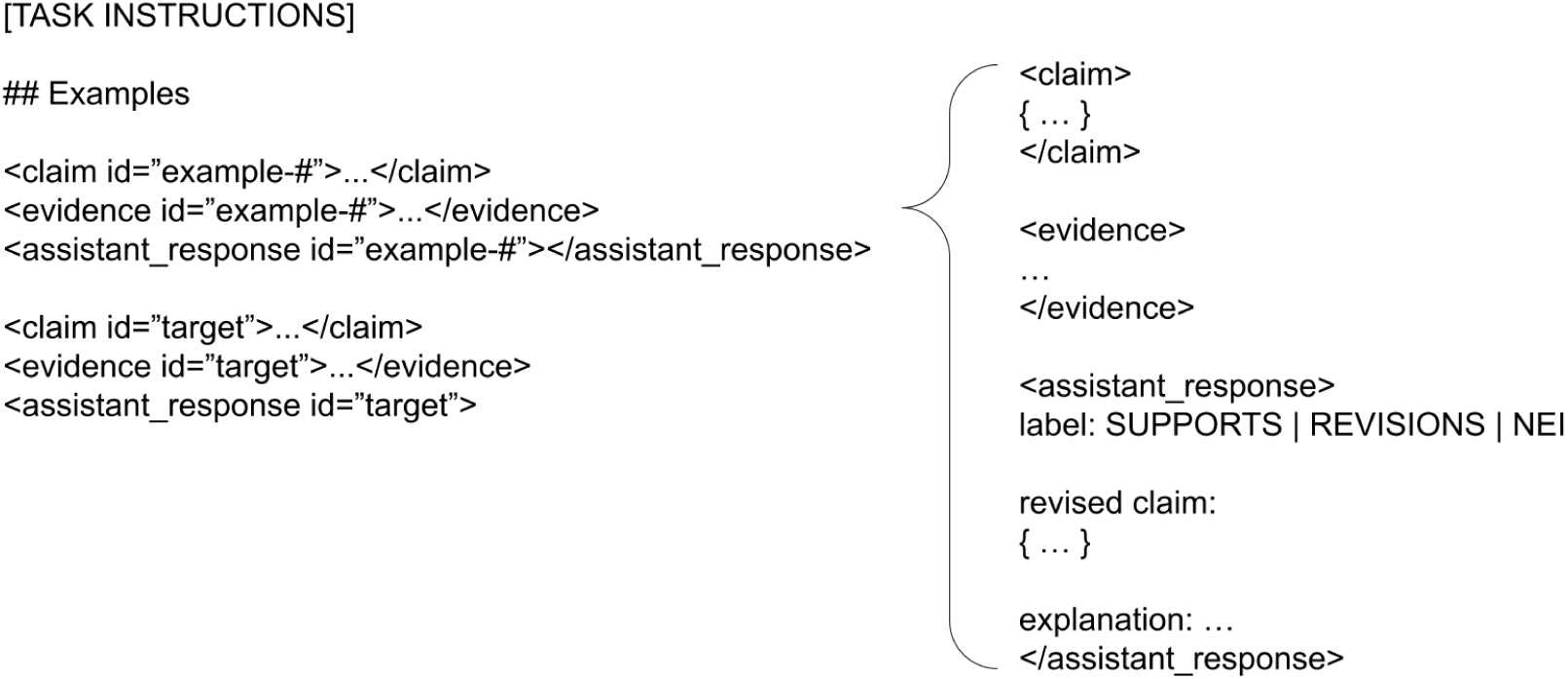
LLM Prompting Format. The prompt begins with an instruction, followed by the claim and evidence pair. Elements specific to a claim/evidence pair are given with placeholders in uppercase surrounded by square brackets and highlighted in blue or yellow. The expected response from the LLM is highlighted in blue. Chat format elements specific to a particular model (in this case Llama3) are highlighted in orange.

We benchmarked a range of language models on CIViC-Fact to assess their ability to verify claims in the CIViC-Fact dataset. Models were given the claim and evidence as input and asked to generate a label and explanation. LLMs that are trained to follow instructions for a variety of tasks use a specific format to denote turns in conversations called chat formats. Chat formats were applied using the transformers^53^ package. Proprietary models were queried using the openai python package. Our experiments included:

● Large Open LLMs including LLaMA-3.1^54^ (70B), Mixtral^55^ (7Bx8E), Gemma 4 (31B); and Llama 4 Scout (17Bx16E), evaluated in few-shot settings (3, 15, or 30 examples)
● A fine-tuned small Open LLM: LLaMA-3.2 (3B)
● Proprietary LLMs: Gemini-2.5^56^; GPT-4.1^57^; and GPT 5.2 evaluated in zero or few-shot settings

### Fine-tuning of local claim-verification models

Local fine-tuning (FT) and evaluation on the CIViC-Fact dataset was completed on up to 8 A6000 GPUs, 2 A100 GPUs, or 8 PRO6000 GPUs depending on availability. Due to resource constraints, each locally run model was given a limit of 400 new tokens for each generation and locally run models were 4-bit quantized. Models were also instructed to keep output explanations to 2-5 sentences. PEFT^58^ and QLoRA^59^ were used in fine-tuning Llama-3.2 3B. The model was trained for completion only^53^ up to 5 epochs with a learning rate of 0.0002 and we used early stopping to avoid overfitting (Supplementary Fig 1). We have made all training code and related configurations available here (https://GitHub.com/creisle/civicfact/tree/master/experiments).

### Historical expert-agreement analysis under the predecessor label formulation

During dataset development, expert agreement was assessed on subsets annotated using the predecessor SUPPORTS/REFUTES/NEI formulation. Annotators achieved a Fleiss’ κ of 0.65 and 77.9% agreement. Because the final benchmark uses SUPPORTS/REVISIONS/NEI, this analysis characterizes the reproducibility of the earlier evidence-stance task and does not directly validate the current REVISIONS category or all reference labels in the final release.

### Expert Human Review of LLM Explanations

To evaluate the explanatory capabilities of LLMs, expert annotators manually reviewed responses generated under the predecessor SUPPORTS/REFUTES/NEI task formulation. Annotators were given a claim–evidence pair and responses from three LLM systems (Llama 3.1 70B, GPT-4.1 and Gemini 2.5). They independently assigned a stance label and explanation under the original three-label system and evaluated the model responses using the categories defined in Table 1. The annotators were asked to both generate a reference label and an explanation for the pair (to be used in future training and automated evaluations) as well as evaluate the responses on the following categories (Table 1). Any cases where the newly generated consensus reference label did not match the expected label were flagged and re-examined to ensure high-quality reference explanations and labels. Because this assessment preceded the introduction of the REVISIONS category, it characterizes explanation quality under the SUPPORTS/REFUTES/NEI formulation and does not directly evaluate explanations generated for the current SUPPORTS/REVISIONS/NEI task.

**Table 1.**
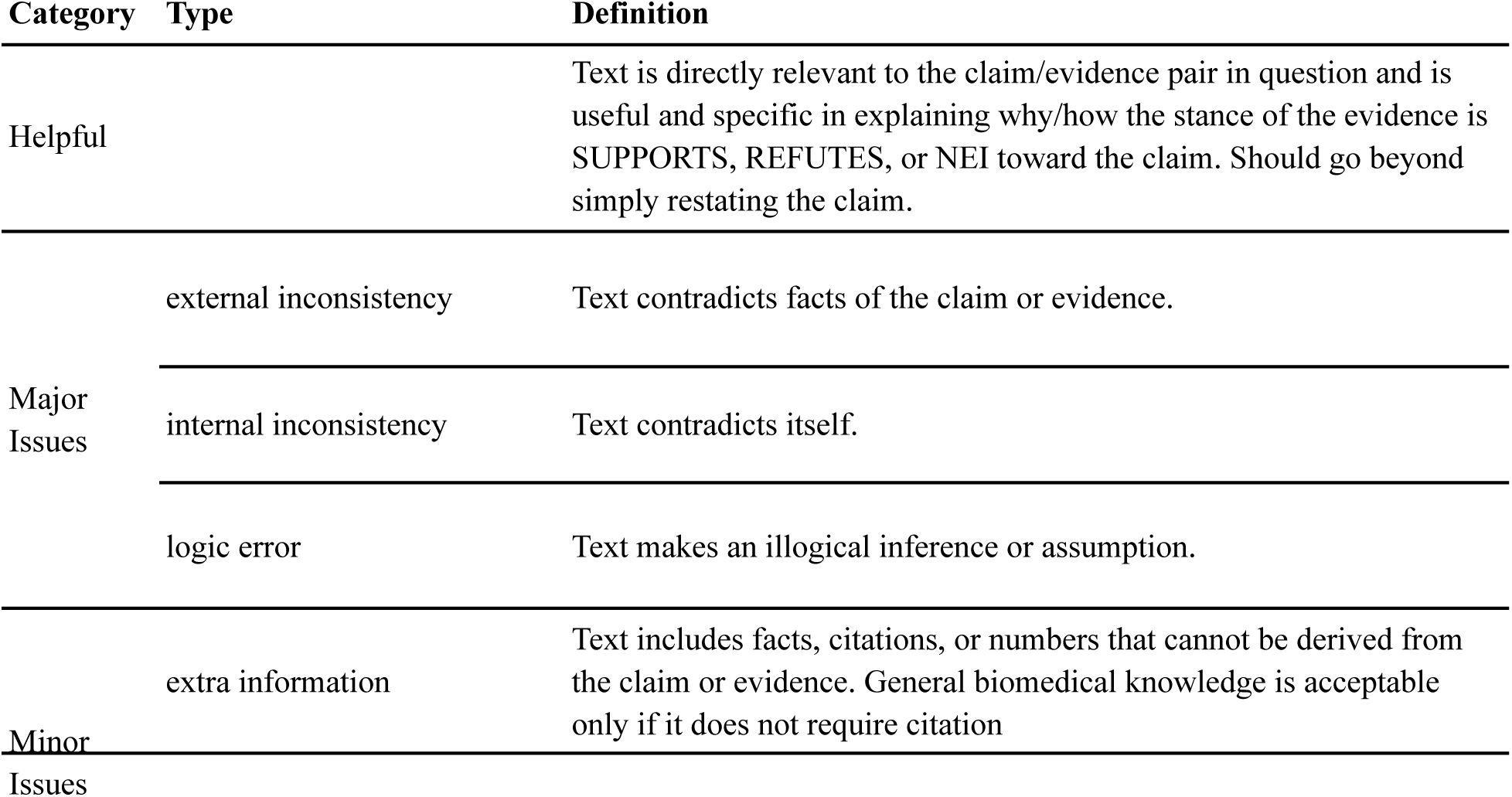
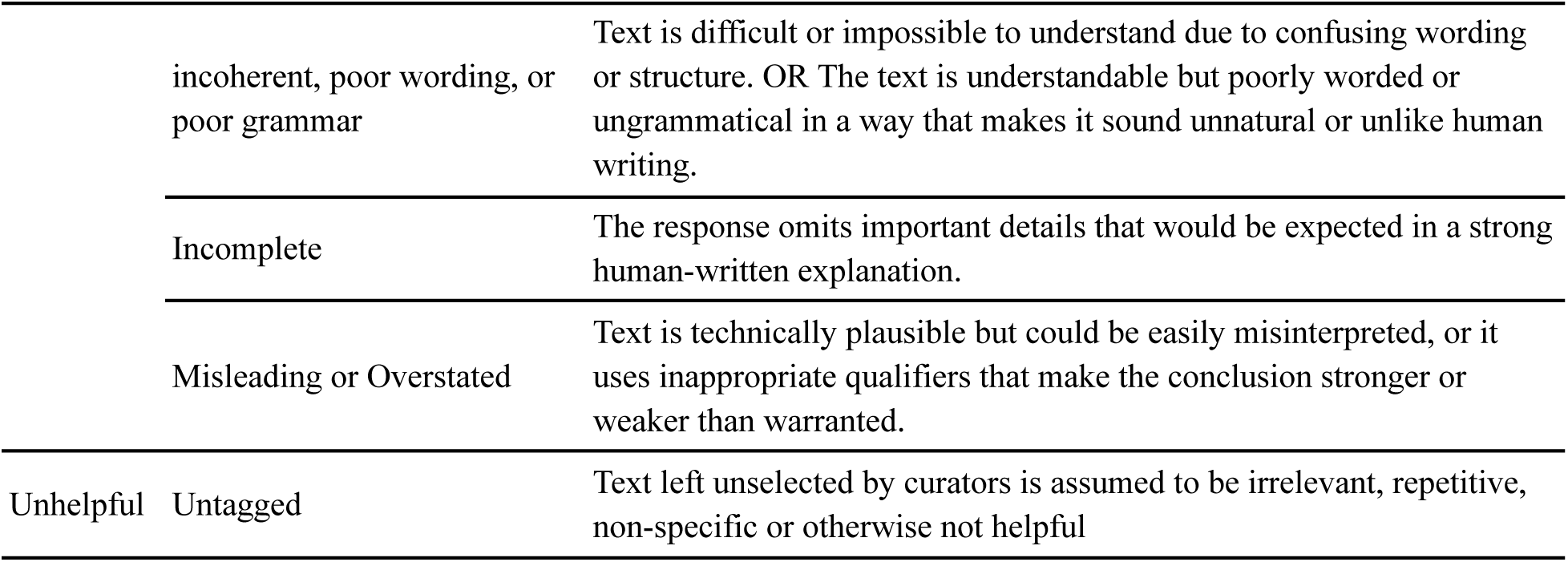
Definitions of tag categories and types used in evaluating LLM response text. The explanation-quality assessment used the predecessor SUPPORTS/REFUTES/NEI formulation; REFUTES was not retrospectively mapped to REVISIONS.

To resolve disagreements from the initial round of tagging, and to capture additional tags such as “*incomplete”* and “*misleading, or overstated”*, each response, along with its initial tags, was re-examined by an additional annotator or collaboratively discussed by the group. We report results for each LLM from this additional analysis. We categorized responses into four groups based on their tagging results: (1) Major Issues, if any major issues were identified; (2) Helpful (Minor Issues), if the response was considered helpful and only minor issues were tagged; (3) Helpful, if the response was helpful with no issues identified; and (4) Unhelpful for all remaining responses.

### Temporal evaluation

In order to apply the CIVIC-Fact pipeline to recent work in CIViC and explore the impact of curation drift we selected CIVIC entries which were submitted or modified after the last build of the CIViC-Fact dataset between 2026-03-03 and 2026-06-09. There were 150 entries, of which 68 were excluded due to missing full-text access, 14 were excluded as they were derived from documents already present in the CIViC-Fact dataset, and 2 were excluded due to non-content related data errors such as being linked to a malformed disease record. Among the remaining 66 entries, a curator evaluated them to determine appropriate reference labels. Any ambiguous cases were discussed as a group until a consensus was reached. The label NEI was given to 24 entries with insufficient evidence that required material outside the main text (images or supplementary material) and 2 where the claim was substantially altered. In the latter case these represent an entry that was re-used to enter a new claim for convenience rather than rejecting the original claim and entering the new claim from scratch. Of the 40 remaining entries with sufficient evidence curators determined 9 to require major revisions that would alter the meaning of the claim, 7 would require minor revisions that alter specificity only and the rest either require no revision or could be trivially revised (Supplementary Figure 2).

## Results

### CIViC-Fact models a full-text curation workflow

CIViC-Fact was designed around the workflow used by CIViC editors to review structured cancer-variant interpretations. Each example begins with a structured CIViC claim and its linked source publication.

Expert annotators identified sentence-level evidence spans within the publication, creating a passage-retrieval task in which a system must surface the portions of the known source document that are relevant to the claim. The retrieved evidence is then used for claim verification, in which a system determines whether the evidence SUPPORTS the claim, indicates that one or more claim fields require REVISIONS, or provides NOT ENOUGH INFORMATION (NEI). When a revision is predicted, the system may also propose a candidate corrected value for curator review. The intended output is therefore decision support for expert curators rather than an autonomous curation decision.

Sentence-level evidence was collected for 1,094 CIViC entries linked to 536 publications. Additional examples were derived from CIViC written descriptions, documented revision histories, controlled modifications of structured claims, and generated negatives designed to represent common curation scenarios. Together, these examples support separate evaluation of passage retrieval and claim verification while preserving the connection between each structured claim and its source evidence (Table 2).

**Table 2.** Quantitative characteristics in each split of CIViC-Fact Claim Verification dataset. The [hard] columns are subsets of their corresponding train, development, and test partitions and contain examples requiring evidence outside the abstract. They are shown separately to characterize these more challenging examples and are not included as additional partitions when calculating totals.

| Task | Characteristic | train | train<br>[hard] | dev | dev<br>[hard] | test | test<br>[hard] | total |
| --- | --- | --- | --- | --- | --- | --- | --- | --- |
| Evidence<br>Retrieval | CIViC entries | 907 | 490 | 99 | 34 | 88 | 35 | 1094 |
|  | passages | 1645 | 800 | 188 | 53 | 172 | 69 | 2005 |
|  | documents | 429 | 164 | 65 | 19 | 42 | 15 | 536 |
|  | documents<br>(full-text only) | 366 | 164 | 57 | 19 | 34 | 15 | 457 |
| Claim<br>Verification | claims | 7418 | 2109 | 845 | 162 | 779 | 199 | 9042 |
|  | documents | 1068 | 214 | 183 | 26 | 138 | 19 | 1389 |
|  | claim/evidence pairs | 9271 | 2432 | 1006 | 182 | 1001 | 231 | 11278 |
|  | NEI | 3222 | 68 | 345 | 1 | 342 | 3 | 3909 |
|  | REVISIONS | 3182 | 1496 | 343 | 114 | 339 | 159 | 3864 |
|  | SUPPORTS | 2867 | 868 | 318 | 67 | 320 | 69 | 3505 |

The benchmark examples were assembled through several provenance routes, including manually selected evidence, existing CIViC content, revision histories and controlled data augmentation. The released labels should therefore be interpreted as provenance-traceable benchmark reference labels rather than uniformly expert-adjudicated ground truth.

### A retrospective audit illustrates potential curation targets

To characterize the types of cases in which an assistive system might be useful, we manually reviewed 100 randomly selected CIViC Evidence Items whose linked publications were available through PubMed Central. The sample included 26 submitted, 70 accepted and four rejected entries. The four rejected entries were excluded from the assessment of existing CIViC content. One additional entry was excluded because the publication contained conflicting evidence and an appropriate revision could not be determined, and another was excluded because it cited an inappropriate non-primary source.

Among the remaining 94 entries, 70 (74.5%) required no revision. Nine (9.6%) were technically consistent with the source evidence but could be represented at a more appropriate level of specificity, such as replacing “PIK3CA expression” with “PIK3CA overexpression.” Ten (10.6%) required updates to their complex molecular profiles, and five (5.3%) contained clear curation errors.

These findings illustrate that AI-assisted review should not be framed solely as error detection. A useful system may also flag scientifically plausible entries that could be updated to reflect greater specificity, newer data-model features or evolving curation guidance. This distinction is particularly relevant for complex molecular profiles, which were introduced after much of the existing CIViC content had already been curated.

The audit also identified an important limitation of a text-only workflow. For two entries, the PubMed Central record contained only an abstract rather than the full article. Eleven additional entries required evidence from supplementary material, figures or both and could not be verified from the main article text alone. The current pipeline should therefore be understood as a proof of concept for main-text evidence retrieval and verification. Cases in which sufficient textual evidence is unavailable should be flagged as outside the supported scope or returned for expert review rather than interpreted as biologically unsupported. Future extensions could incorporate supplementary files, tables, figures and other multimodal evidence.

### Abstracts are insufficient for realistic cancer variant claim verification

The development partition of the CIViC-Fact retrieval dataset contains 99 CIViC entries with one or more sets of annotated sentence-level passage retrieval evidence for each. Of these 87 are linked to full-text documents and 12 are linked only to abstracts. Of the remaining 87, 2 cannot be validated by text alone and require images or supplementary material. Among the 85 text-verifiable development entries with full-text access, 24 (28.2%) could be fully validated from the abstract alone, 34 (40.0%) required evidence entirely outside the abstract, and 24 (28.2%) were only partially supported by the abstract. This means that nearly 70% of CIViC entries in this subset cannot be validated by the abstract alone (Supplementary Fig 3). We call the subset that requires evidence entirely outside the abstract (dev[hard]) and we report these results separately. This is of particular importance as most biomedical NLP work has focused on short documents or abstracts^33,44,60,61^ rather than full text and inter-rather than intra-document retrieval.

Even biomedical datasets focusing on long documents such as LongHealth^62^ (133 documents, average of 6.39k characters per document) still have documents that are considerably shorter than the average scientific article. For example, there are 457 full-text scientific articles in the full CIViC-Fact dataset which average 32.3k characters per document (Fig 6a), more than 5x longer than the documents in the LongHealth^62^ dataset. This presents a significant challenge to retrieval models when evidence supporting a claim could be anywhere in the document.

**Figure 6.**
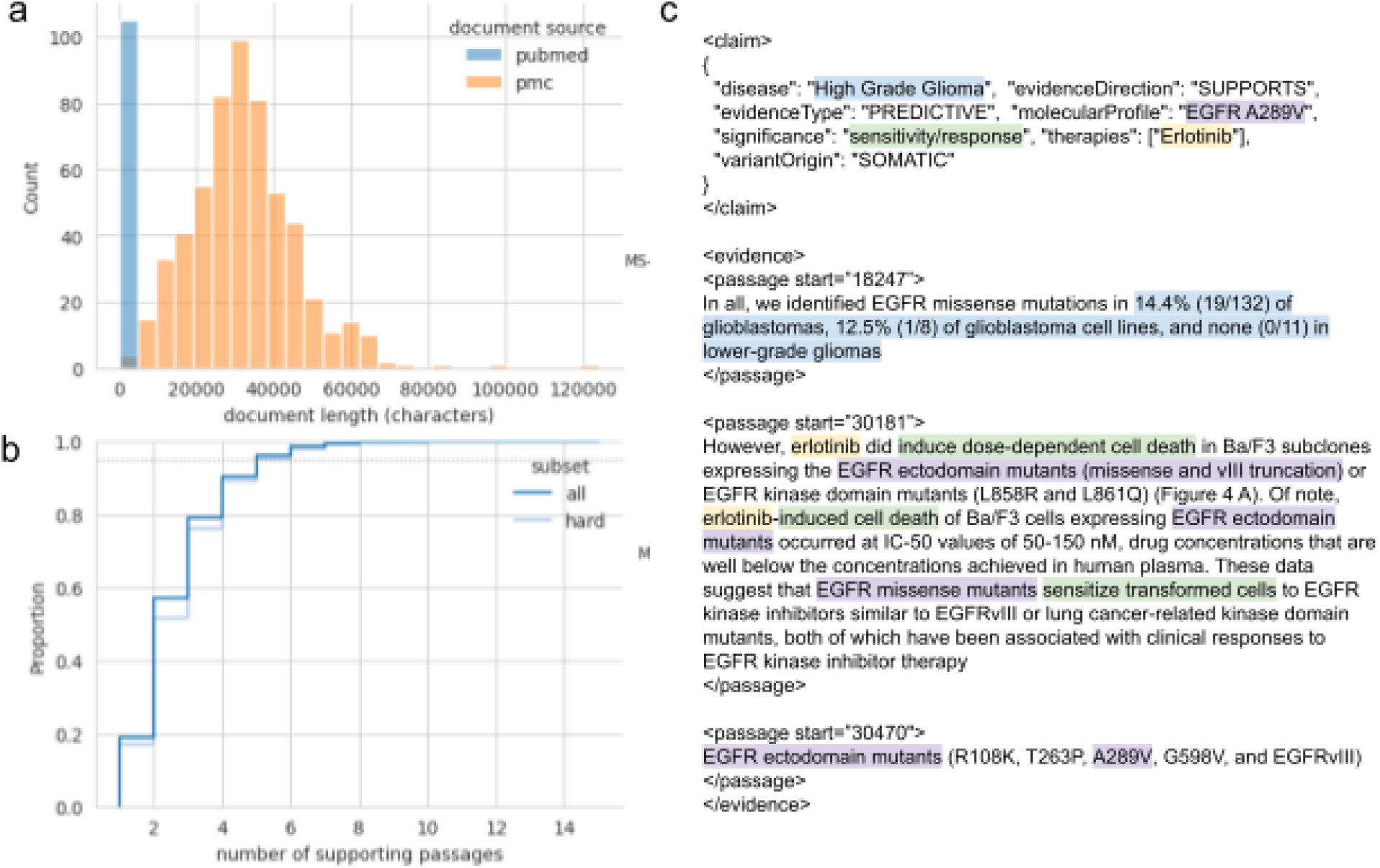
a. Document length by number of characters for documents in the CIViC-Fact dataset. b. Number of passages overlapping manually annotated evidence passages for each example in the retrieval portion of the CIViC-Fact dataset. The 95th percentile is given with a horizontal dashed black line. c. Impact of number of passages retrieved (k-value) on mean average precision. c. A CIViC-Fact example requires 3 distinct passages from distant parts of the source document. The first passage identifies the claims relevance to the curated disease “High Grade Glioma” whereas the second passage describes the preclinical experiments detailed the response of this variant to treatment with erlotinib. Finally the third passage is required to disambiguate which particular variants were studied in passage 2.

Furthermore, CIViC-Fact contains examples with complex claims where validation requires passages from multiple (between 1-15 passages, 2 on average: Fig 6b) sometimes distant parts of the source document. For downstream evaluations, we chose to limit the retrieval to the top 5 passages as this covers the required selections for 95% of the dataset (Fig 6b). Generally, claims in the hard-retrieval dataset contain more logical “hops”^37,63,64^ than those that can be found in the abstract since the abstract represents a summarization of the article (Fig 6c).

### Automated passage retrieval can surface evidence for expert review

To fact-check CIViC claims at scale, we incorporated automated passage retrieval to identify the most relevant text segments from the source paper, rather than relying on manual curation. Using the linked source paper for each claim, embedding models ranked candidate passages by relevance, with the top results provided as evidence for claim verification (Fig 3). We compared general and biomedical-specific models (Specter2^41^, MedCPT^44^, Qwen 3 reranker^50^, BGE reranker^45^) against the BM25^51^ baseline. We include the first k passages (head) as an additional baseline. We measured the performance of passage retrieval models using the mean average precision (mAP) which scores rankings of passages based on the position of relevant passages. A score of 1 indicates that all of the top-k passages retrieved are relevant.

The best performing model was also the largest retrieval model used, the Qwen 3 reranker (8B)^50^. Qwen 3 reranker [8B] outperformed the other non-fined-tuned passage retrieval models. We also fine-tuned some of the smaller passage retrieval models on the CIViC-Fact dataset. These fine-tuned models performed similarly to Qwen3 and even demonstrated minor improvement on the hard subset. Fine-tuned MedCPT achieved 0.51 on the hard subset whereas Qwen 3 only achieved 0.48 (Table 3). We test both the fine-tuned MedCPT and the Qwen 3 reranker [8B] (hereafter referred to simply as Qwen 3) model in downstream tasks.

**Table 3.** Retrieval accuracy (k=5) as measured by mean average precision (mAP) using the best-performing query-template for each model. The head baseline takes the first 5 passages. The highest mAP in each column is bolded. The baseline models are given first followed by the models evaluated in decreasing order of mAP based on the full development partition.

| model | dev | dev[hard] | test | test[hard] | query template |
| --- | --- | --- | --- | --- | --- |
| head | **0.53** | 0.08 | 0.46 | 0.04 |  |
| BM25 | 0.47 | 0.41 | 0.47 | 0.36 | NL |
| Qwen 3 [8B] | **0.53** | 0.48 | **0.58** | **0.51** | col is value; |
| MedCPT (FT) | 0.52 | **0.51** | 0.49 | 0.49 |  |
| MS-MARCO-MiniLM | 0.50 | 0.36 | 0.44 | 0.33 | NL |
| ModernBERT (FT) | 0.50 | 0.46 | 0.49 | 0.41 |  |
| Qwen 3 [4B] | 0.49 | 0.38 | 0.49 | 0.42 | col is value; |
| Qwen 3 [0.6B] | 0.47 | 0.40 | 0.45 | 0.33 | col is value; |
| MedCPT | 0.46 | 0.35 | 0.46 | 0.34 | NL |
| BGE | 0.44 | 0.38 | 0.49 | 0.43 | NL |
| ModernBERT | 0.44 | 0.42 | 0.38 | 0.33 | NL |

Retrieval evaluation should be interpreted carefully. Mean average precision provides a useful ranking metric, but the manually annotated evidence in CIViC-Fact is not necessarily exhaustive. And while mean average precision (mAP) provides a measure of how relevant the retrieved passages are, it does not indicate whether those passages are sufficient to verify a claim. Scientific articles often contain redundant or overlapping statements, and a passage not originally annotated by a curator may still be sufficient to verify a claim. For this reason, the manual review of retrieval outputs on newly submitted CIViC entries is an important complement to automated retrieval metrics. It provides a more realistic estimate of whether retrieved passages are sufficient for downstream curation, rather than simply measuring whether they reproduce the exact original evidence selection. Some passages retrieved by the model but not annotated in the dataset may in fact be relevant, false negatives that could serve as valid substitutes for the annotated passages. Therefore, in order to better determine an accurate estimate of the true retrieval accuracy, we refined this estimate through manual review.

We applied fact-checking, and therefore retrieval, to newly submitted CIViC entries collected after the cutoff date for the CIViC-Fact dataset, spanning 3 March to 9 June 2026. Among this subset there were 42 entries for which full-text was available and the claim did not require supplementary material or images for verification. 2 were excluded as the revised claim was substantially different from the original claim and they were given the reference label of NEI (Supplementary Fig 2). Consistent with the mAP results, both the fine-tuned MedCPT model and the 8B Qwen 3 reranker performed similarly retrieving appropriate content for 37 of the remaining 40 entries (92.5%). The fine-tuned MedCPT model however was more resource efficient as it is a much smaller non-generative model and could be run on a single GPU. It was therefore used in downstream claim-verification experiments.

### Evaluating fine-tuned and few-shot models for claim verification and candidate revision generation

Claim verification is defined as a model’s ability to correctly label the stance of evidence toward a claim when given both the claim and evidence as input. To get a measure of language models reasoning performance when the expected evidence was provided, we evaluated language models on the CIViC-Fact dataset with the expert-selected evidence collected from hypothes.is or simulated examples (see methods). Prompt and in-context-learning configurations were optimized on the development partition before final evaluation. Full optimization procedures and selected configurations are described in the Supplementary Methods.

On the held-out test partition, Gemma 4 achieved 82% accuracy, GPT-5.2 achieved 79%, and the fine-tuned Llama 3.2 3B model achieved 77%. Models were evaluated on both the full partition as well as the hard-retrieval subset for both development and test partitions (Fig 7a). The hard retrieval subset contains only examples where the target passages are not found in the abstract. In general, models performed worse (approximately a 10% drop in accuracy for all models) on the hard-retrieval dataset despite being given the required passages. This indicates that the difficulty in this subset is not only in its retrieval but also in its reasoning.

**Fig 7.**
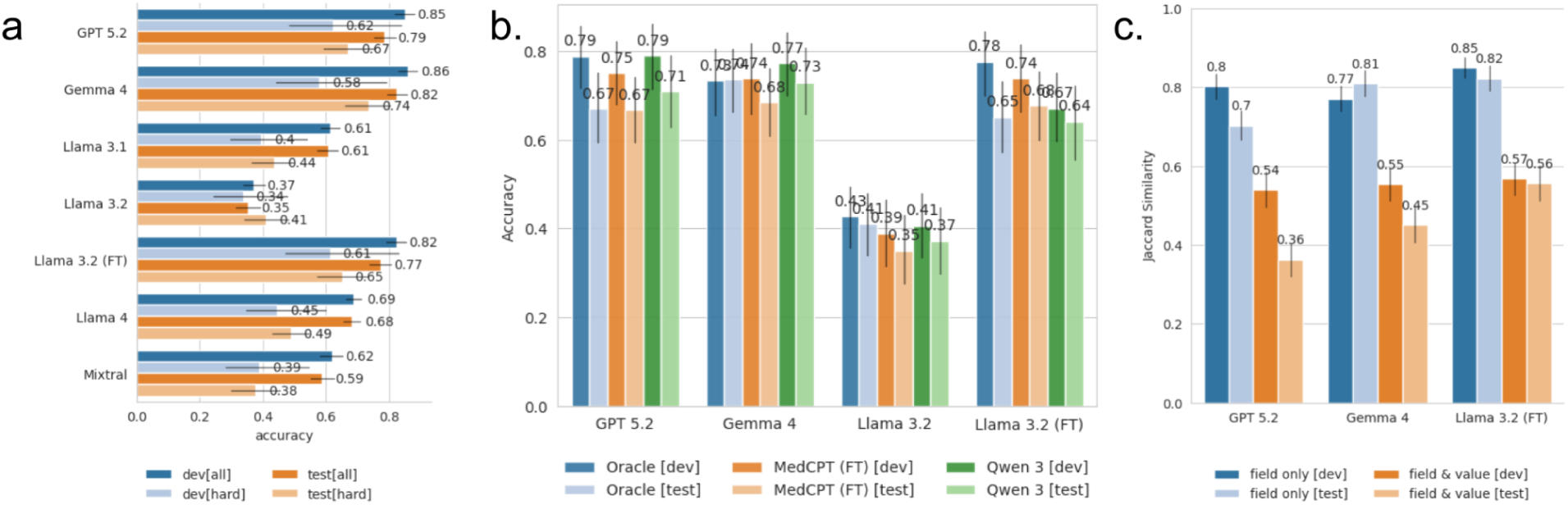
(a). Label prediction accuracy on the development and test partitions of the CIViC-Fact dataset. Accuracy is reported per label or macro-averaged (average). We exclude NEI from the macro-average of the hard subset as it contains insufficient examples for this label. (b). Accuracy of the top-3 (GPT 5.2; Gemma 4; fine-tuned Llama 3.2) performing models on the hard subset of the development and test partitions including with and without (Oracle) the evidence retrieval step. We report retrieval results for the fine-tuned MedCPT model, the 8B Qwen 3 reranker. The oracle is included as a positive control. It is the first 5 passages that overlap manually annotated evidence. These may include repetitive content and fail to include key content as they are selected from all evidence annotated for a particular CIViC entry and not independent sets of evidence. (c) Model accuracy in predicting revisions on the development and test partitions of the CIViC-Fact dataset.

In practice, the pipeline would be composed of both passage retrieval and claim verification components. We tested the top three performing models, Gemma 4, GPT 5.2, and the fine-tuned Llama 3.2, on the combined retrieval and verification task. We also evaluated the classification accuracy when the claim verification model was given evidence from a passage retrieval model, rather than the evidence manually selected by the expert curators (Human). We include an “Oracle” retrieval estimate by selecting k passages from the document, first selecting any annotated for the claim by a human curator and then based on their position in the document, preferring passages earlier rather than later. This should be similar to the Human selected evidence but with additional distractor passages since it will include non-relevant evidence when there are less than 5 human selections. Furthermore the passages will be in 1000 character chunks rather than following sentence boundaries. The fine-tuned model achieved 68% accuracy on the test data, a 9% drop over that achieved with the human selected evidence. Results for the few-shot models were similar to the classification accuracy we saw with the Human-selected evidence: GPT 5.2 and Gemma 4 did not show any significant change in accuracy due to the possible distractors passages (Fig 7b).

In addition to checking the label accuracy, we also evaluate the accuracy of the suggested revision that the model produces by checking all true revisions against their expected revision. However, as multiple claims can be derived from the same article it is still possible the models are producing valid, if unexpected, revisions. Therefore, we also compare the suggested revision to all known claims from the same document. Our fine-tuned language model achieves the highest accuracy in picking the correct fields to revise, however when we measure the accuracy of both the field revised and the value it was revised to, all 3 models perform poorly (Fig 7c). Exact matches are still low, leaving room for improvement. For example, some models produce non-preferred, or deprecated, disease ontology terms where we expect the preferred term. This is still semantically plausible but not an acceptable exact match. Future work will explore the inclusion of additional ontology information in the context to further improve outcomes.

### Temporal evaluation reveals curation drift and limits of static fine-tuning

We evaluated the three final claim-verification systems on the temporally held-out cohort described above (Supplementary Fig. 2). For each claim, the five highest-ranked passages were retrieved independently using either the fine-tuned MedCPT retriever or Qwen3-8B reranker, according to the retrieval configuration that performed best on the development partition. Qwen3-8B retrieval was used for the few-shot models, whereas the fine-tuned MedCPT retriever was paired with the fine-tuned Llama 3.2 claim-verification model. In parallel, a CIViC curator manually reviewed each claim and its source evidence to assign a reference label. Challenging or ambiguous cases were discussed by the CIViC curation team until consensus was reached, and model predictions were compared with these curator-assigned reference decisions.

While all systems performed worse on this heterogeneous post-cutoff cohort than on the retrospective CIViC-Fact evaluation set, performance was considerably lower for the fine-tuned model. Notably, the post-cutoff cohort was enriched for particular curation activities and therefore was not representative of the current CIViC knowledgebase as a whole. It also contained a disproportionate number of claims for which verification depended on information reported only in images or supplementary material (24 of 66). Such cases are comparatively uncommon in CIViC and, consequently, were sparsely represented in the CIViC-Fact training data. We therefore stratified performance according to whether the information required to verify a claim was available in the retrieved main-text evidence (‘sufficient evidence’) or could not be recovered from the available passages (‘insufficient evidence’).

Among claims with sufficient evidence, the two few-shot systems performed similarly, with Gemma and GPT achieving accuracies of 64%, compared with 45% for the fine-tuned system. A substantially larger difference emerged for claims with insufficient evidence. GPT and Gemma correctly identified 75% and 88% respectively of these cases, whereas the fine-tuned system did not correctly identify any. Thus, the principal performance gap on the post-cutoff cohort arose not from verification of claims for which the relevant information was available, but from recognizing when the retrieved evidence was insufficient to support a reliable curation decision. Previous work has explored evidence insufficiency as a standalone task^38^. A future avenue we might explore with fine-tuning would be to split the current verification task into two steps. One to determine the sufficiency of the evidence and a second step for claim verification when evidence is sufficient. This would allow us to train with more balanced and targeted datasets as training all 3 labels at once results in a significantly larger sample space for the NEI class compared to the REVISIONS class and the REVISIONS class compared to the SUPPORTS class. It is also likely that some of the performance lost for the fine-tuned model can be explained by changes in the curation process. The post-cutoff cohort represents current guidelines and recommendations in curation whereas the training data is more static and fixed on best practices at the time it was collected.

This distinction is particularly relevant for CIViC because curation represents a dynamic process in which annotation guidance continues to evolve. During development of CIViC-Fact, several curation guidelines were refined in response to ambiguities identified during expert review, including cases in which experienced editors initially reached discordant interpretations. One example concerned assignment of disease terms for preclinical evidence. A disease term may reflect either the biological model used experimentally—for example, assigning ‘breast cancer’ to evidence generated in MCF-7 cells—or the broader disease context intended by the study authors. Thus, for a variant identified across several tumour types and subsequently evaluated for drug sensitivity in selected cell lines, a model-centric interpretation would assign disease according to the tissue of origin of the experimental cell line, whereas an author-centric interpretation could favour a broader term such as ‘cancer’ when the study was explicitly pan-cancer in scope.

A second area of evolving guidance concerned representation of multiple variants within complex molecular profiles. Complex molecular profiles were introduced into CIViC in January 2023, after much of the existing knowledgebase had already been curated, and some earlier entries therefore pre-date this representation framework. During the present curation effort, guidance was further refined regarding whether a secondary mutation should be incorporated into the molecular profile. In cases involving a primary driver alteration together with a secondary resistance mutation, ambiguity can arise when the driver alteration is not represented in the experimental control. The resulting guidance favoured inclusion of the driver event unless multiple distinct driver events were described, in which case omission of the driver could be appropriate.

The fine-tuned model captured some previously established curation conventions, including the use of complex molecular profiles, consistent with these patterns being represented in its training data. However, a fixed fine-tuned model cannot directly incorporate guideline changes introduced after construction of the training set. This limitation may have contributed to its reduced performance on the post-cutoff cohort, although differences in cohort composition and evidence availability are also likely to have contributed. By contrast, few-shot prompting permits updated curation principles to be introduced through a small number of revised examples. These results therefore suggest that, for knowledgebases in which curation standards continue to evolve, few-shot approaches may offer greater flexibility than relying exclusively on model fine-tuning.

## Discussion

CIViC-Fact provides a proof of concept for using artificial intelligence to supplement expert review of structured cancer-variant interpretations. It models three linked components of curation: retrieving relevant passages from a known source publication, determining whether those passages support a structured claim or indicate that it requires revision, and proposing candidate corrections for curator consideration. Domain experts manually identified sentence-level evidence from the source literature, while claim-verification labels were assembled from existing CIViC records, documented revision histories and controlled data augmentation. CIViC-Fact is therefore an expert-curated, provenance-rich framework for evaluating AI-assisted curation.

Its full-text design is central to this contribution. CIViC-Fact contains 11,278 claim–evidence pairs from 1,094 CIViC entries and 536 publications, including revision examples and generated negatives intended to represent common curation problems. Fewer than one-third of the evaluated text-verifiable development entries could be fully assessed from the abstract alone, and some required evidence from multiple distant passages. Because relevant evidence may appear anywhere in a long article, abstract-only evaluations may overestimate model readiness for biomedical curation by omitting cases that require retrieval and synthesis across methods, results, tables and other non-abstract content.

The retrieval results support the feasibility of automatically surfacing candidate evidence for expert review. Among text-verifiable entries in the post-cutoff cohort, appropriate evidence was retrieved for 37 of 40 cases. This supports a human-in-the-loop workflow in which retrieval models direct curators to relevant portions of a linked publication, language models assess the relationship between the claim and evidence, and expert editors review the evidence, label and proposed revision before accepting any change.

At the same time, main-article text was not sufficient for every case. In the retrospective audit, 11 of 94 evaluable entries required evidence from supplementary material, figures or both, and two additional PubMed Central records contained only abstracts rather than full articles. In the post-cutoff cohort, 24 of 66 entries required supplementary or image-based evidence. The higher proportion in the latter cohort likely reflects its enrichment for particular curation activities and should not be interpreted as representative of CIViC overall. Together, these analyses show that the coverage of a main-text-only system depends on cohort composition. Cases lacking sufficient main-text evidence should therefore be identified as outside the supported scope of the current workflow rather than interpreted as evidence that the biological claim is unsupported.

CIViC-Fact may also support retrospective knowledgebase review. In the audit of 94 non-rejected entries, 70 required no revision, nine were technically correct but could be represented with greater specificity, ten required updates to a complex molecular profile and five contained clear curation errors. These findings show that entries may warrant reassessment because of changes in the CIViC data model or curation guidance, not only because of an original curator error. Early operational use provides qualitative support for this application: the workflow surfaced several existing entries that were subsequently found to require correction or reassessment. Because CIViC entries are not routinely re-reviewed and are generally revisited only when independently flagged, AI-assisted screening provides an additional mechanism for bringing potentially problematic legacy content to expert attention.

The post-cutoff cohort further highlights the challenge of applying static models to a living knowledgebase. As CIViC guidance and its data model evolve, models trained on historical examples may reproduce outdated conventions. Because this cohort was enriched for specific curation activities, differences between models should be interpreted as illustrative rather than representative of CIViC overall. Nevertheless, the reduced performance of the fine-tuned model relative to few-shot systems suggests that maintaining performance in a changing curation environment requires mechanisms for incorporating updated guidance. Updating a small set of in-context examples offers a substantially lower-effort adaptation strategy than constructing a new training dataset, retraining and revalidating the model.

The model comparisons also distinguish benchmark performance from long-term curation utility. Fine-tuning a small local model substantially improved agreement with CIViC-Fact reference labels on the static benchmark, showing that targeted domain adaptation remains valuable when training and evaluation data are closely matched. Local models may also offer advantages in governance, availability, cost and institutional control. However, the fine-tuned model was less robust on heterogeneous newer submissions, particularly when the available evidence was insufficient or curation conventions had changed. Fine-tuning should therefore be viewed as one deployment option rather than a complete maintenance strategy for a changing knowledgebase.

The revision generation results reinforce the need for continued human oversight. Correctly assigning a REVISIONS label is insufficient if the proposed edit is unusable. The fine-tuned model performed well at identifying which fields required attention, but exact agreement on both the revised field and proposed value remained limited. Some outputs were conceptually plausible but used deprecated or non-preferred ontology terms. These findings identify clear opportunities for improvement, including providing models with current ontology terms, hierarchical relationships and curation guidance at inference time. Incorporating this context may improve the precision and practical usefulness of proposed revisions while preserving expert review as the final decision point.

The historical explanation analysis provides a further reason for caution. Under the predecessor SUPPORTS/REFUTES/NEI formulation, expert reviewers found that model explanations could contain useful biomedical reasoning but were not consistently reliable. Some responses identified subtle inconsistencies, whereas others introduced unsupported details, factual errors or invalid inferences. Because this analysis used an earlier label formulation, model set and prompting configuration, it should not be interpreted as a current comparison of explanations for the REVISIONS task. Its broader lesson remains relevant: fluent rationales cannot be assumed to be evidence-grounded, and model evaluation should consider explanation quality as well as label agreement.

A secondary contribution of CIViC-Fact is its use as a reproducible benchmarking infrastructure. Models, hosted services and retrieval systems change rapidly, making conclusions about any single system time-dependent. CIViC-Fact can be rerun as models and pipeline components change, allowing performance to be compared against consistent task definitions and evidence provenance. Because retrieval, claim verification and revision generation are evaluated separately, individual components can be updated without rebuilding the entire workflow. This benchmarking role complements rather than replaces prospective evaluation in expert curation practice.

Overall, CIViC-Fact demonstrates the feasibility of using NLP-based tools to supplement expert biomedical curation. Retrieval models can surface candidate evidence, language models can classify claim/evidence relationships and propose revisions, and expert curators can review the outputs before approving any change. Early operational experience also suggests that the workflow can identify legacy entries that would not ordinarily be revisited without an independent flag. CIViC-Fact’s primary contribution is therefore an expert-curated, full-text proof-of-concept framework for human-in-the-loop cancer variant curation, with the additional benefit of supporting reproducible comparison of retrieval and verification systems as models and curation standards evolve.

## Limitations and Future Work

While our system shows promising results, several limitations remain. First, we only locally tested smaller quantized models, while this is reasonable given that likely any centre would need to account for resource constraints it is not a strictly fair comparison of the open LLM models capabilities.

CIViC-Fact also clarifies the limits of text-only full-text verification. Full-text main-article evidence is a major improvement over abstract-only evaluation, but it is not the final form of biomedical fact-checking. In the manuscript’s manual estimate, approximately 12% of reviewed CIViC entries could not be verified from main text alone because they required supplemental material, images, or both. In the newer-submission retrieval analysis, a substantial number of entries were excluded because the relevant evidence was in supplemental content. Future versions of this benchmark should therefore incorporate supplementary files, tables, figures, and multimodal evidence once sufficient examples are available. Until supplementary and multimodal evidence are incorporated, NEI should be interpreted as indicating that the evidence available to the text-only verifier is insufficient to assess the claim. In some cases, the relevant support may be contained in supplementary materials, figures or other evidence outside the current workflow. NEI reflects insufficient accessible evidence, not a judgement that the biological claim is unsupported.

Both the inter-annotator agreement analysis and the expert assessment of LLM explanations were conducted using the predecessor SUPPORTS/REFUTES/NEI formulation rather than the current SUPPORTS/REVISIONS/NEI task. Consequently, these analyses do not directly validate the current REVISIONS reference labels or evaluate whether models can reliably explain and propose appropriate revisions. The explanation assessment also evaluated an earlier set of models and prompting configurations, including zero-shot responses from Gemini 2.5 and GPT-4.1. Because repeating this specialist review was not feasible, the reported explanation results should be interpreted as evidence that fluent model rationales may contain substantive errors, rather than as a current comparison of model explanation quality.

We envision future extensions of this work that integrate additional annotator feedback, incorporate uncertainty quantification in model predictions, and expand the scope of verifiable claims to include multi-modal evidence and supplementary materials. With continued development, we believe this approach can help foster more trustworthy, evidence-backed cancer knowledgebases.

## Data and Code Availability

| Model Name | Huggingface ID or Model Version | Model Size (B) |
| --- | --- | --- |
| BERT | [google-bert/bert-large-uncased](https://huggingface.co/google-bert/bert-large-uncased) | 0.3 |
| BGE | [BAAI/bge-reranker-large](https://huggingface.co/BAAI/bge-reranker-large) | 0.6 |
| BioMedBERT | [microsoft/BiomedNLP-BiomedBERT-base-uncased-abstract-fulltext](https://huggingface.co/microsoft/BiomedNLP-BiomedBERT-base-uncased-abstract-fulltext) | 0.11 |
| BioMistral | [BioMistral/BioMistral-7B-DARE](https://huggingface.co/BioMistral/BioMistral-7B-DARE) |  |
| Gemini 2.5 | gemini-2.5-pro-preview-03-25 | not given |
| GPT 4.1 | gpt-4.1-2025-04-14 | not given |
| GPT 5.2 | OpenAI GPT5.2 | not given |
| Llama 3.1 | [meta-llama/Llama-3.1-70B-Instruct](https://huggingface.co/meta-llama/Llama-3.1-70B-Instruct) | 70 |
| Llama 4 | [meta-llama/Llama-4-Scout-17B-16E-Instruct](https://huggingface.co/meta-llama/Llama-4-Scout-17B-16E-Instruct) | 272 |
| MedCPT | [ncbi/MedCPT-Cross-Encoder](https://huggingface.co/ncbi/MedCPT-Cross-Encoder) | 0.33 |
| Mixtral | [mistralai/Mixtral-8x7B-Instruct-v0.1](https://huggingface.co/mistralai/Mixtral-8x7B-Instruct-v0.1) | 56 |
| ModernBERT | [tomaarsen/reranker-ModernBERT-large-gooaq-bce](https://github.com/tomaarsen/reranker-ModernBERT-large-gooaq-bce) | 0.4 |
| MS-MARCO-MiniLM | [cross-encoder/ms-marco-MiniLM-L12-v2](https://github.com/cross-encoder/ms-marco-MiniLM-L12-v2) | 0.334 |
| Qwen 3 reranker [0.6B] | [Qwen/Qwen3-Reranker-0.6B](https://github.com/QwenLM/Qwen3-Reranker-0.6B) | 0.6 |
| Qwen 3 reranker [4B] | Qwen/Qwen3-Reranker-4B | 4 |
| Qwen 3 | Qwen/Qwen3-Reranker-8B | 8 |
| Qwen 3.5 [27B] | [Qwen/Qwen3.5-27B](https://github.com/QwenLM/Qwen3.5-27B) | 27 |
| Qwen 3.5 [9B] | [Qwen/Qwen3.5-9B](https://github.com/QwenLM/Qwen3.5-9B) | 9 |
| Specter | [allenai/specter](https://github.com/allenai/specter) |  |
| Specter2 | [allenai/specter2_base](https://github.com/allenai/specter2_base) |  |
| Gemma 4 | [google/gemma-4-31B-it](https://github.com/google/gemma-4-31B-it) | 31 |

Versions of the CIViC-Fact dataset can be downloaded from GitHub (https://GitHub.com/creisle/civicfact) along with unprocessed data from manual reviews. A frozen download of the CIViC *Evidence Items* used in this analysis is also included for reproducibility. Code for building the dataset and running language model experiments can be found on GitHub (https://GitHub.com/creisle/civicfact) as can the forked version of doccano^65^ with additional feature support used for the manual review tasks (https://GitHub.com/creisle/doccano).

## Use of Large Language Models (LLMs) in the writing process

Generative artificial intelligence tools (ChatGPT, OpenAI, San Francisco, CA) were used to refine the clarity and readability of the manuscript text. The tool was not used for data analysis or interpretation. All outputs were carefully reviewed and edited by the authors to ensure accuracy and appropriateness, and the final manuscript reflects the authors’ own work and conclusions.

## Supplementary Methods

### Fine-tuning passage retrieval models

We used the sentence-transformers^46,66^ package to train and evaluate passage retrieval models. The CIViC-Fact data most closely resembles the “anchor-positive” pairs. Therefore we used the builtin functionality of the sentence transformers to mine “hard negatives” both for training and evaluation. We chose the Cached Multiple Negatives Ranking loss function in order to effectively include more in-batch negatives. Training was completed on a A6000 GPU with an effective batch size of 64 and a learning rate of 0.00002. Models were trained up to 5 epochs using early stopping to avoid overfitting. Normalized Discounted Cumulative Gain^67^ at rank 10 (ndcg@10) was used to evaluate model performance.

### Prompt optimization and model configuration

We evaluated the effects of prompt formulation and in-context learning (ICL) on model performance. Specifically, we varied the system prompt, user prompt format, and number of ICL examples provided to the model (3, 15, or 30 examples). We also generated natural-language explanations for all claim–evidence pairs in the training partition using GPT-5.2 and assessed whether including these explanations as ICL examples or during fine-tuning improved model responses.

Because evaluating large locally hosted models and proprietary models across all configurations is computationally and financially intensive, configuration optimization was performed on randomly selected subsets of the development partition comprising 10% (system prompt) or 20% (all other conditions) of examples depending on the task. Only the final selected configuration for each model was evaluated on the full dataset. Configuration parameters were explored individually, and final configurations were selected under the assumption that parameter effects were approximately additive or, at minimum, non-interfering. Although an exhaustive search over all parameter combinations could potentially yield improved performance, such an analysis was not feasible within available resources.

### System-prompt conditions

We evaluated three system-prompt conditions. In the first, no system prompt was provided. In the second, a general-purpose “ChatGPT-like” system prompt was used. In the third, termed the CIViC-editor prompt, GPT-5.5 was used in agent mode through the web interface with extended reasoning enabled to draft a task-specific system prompt after reviewing CIViC curator documentation. The resulting prompt was manually reviewed and corrected for accuracy before use.

### User-prompt formats

We tested four user-prompt formats: simple, full-text, evidence, and passages. The full-text prompt framed the task as combined passage retrieval and claim verification, in which the model was provided with the complete article and asked to identify relevant passages and classify the stance of the evidence. Owing to context-length constraints, this format was evaluated only in zero-shot settings and only for models supporting long-context inputs. The evidence and passages formats differed only in the representation of the evidence text. In the evidence format, passages from different parts of the article were concatenated with ellipses. In the passages format, each passage was enclosed within ‘<PASSAGE>’ XML-style tags and annotated with its start and end character positions in the source article, providing the model with information about the relative locations of the evidence within the article.

### In-Context Example Selection

For non-fine-tuned models, we evaluated ICL using 3, 15, or 30 examples. In all cases, when performance differences between configurations were negligible, the simpler configuration was selected for subsequent evaluation. In-context examples were randomly selected but balanced across label types. Therefore when 3 examples were given the model received one of each: SUPPORTS, REVISIONS, and NEI.

### Explanation-augmented prompting/fine-tuning

Explanations were generated for the training data in the CIViC-Fact dataset using GPT5.2. We then tested whether including these explanations in the examples provided to the model affected accuracy. d. Models were tested with 3, 15, or 30 examples.

### Expert Human Review of the CIViC-Fact Claim Verification Dataset

Expert review was conducted during earlier iterations of CIViC-Fact to assess evidence-stance judgments and the quality of generated content. These annotation rounds used the SUPPORTS/REFUTES/NEI formulation, whereas the final benchmark uses SUPPORTS/REVISIONS/NEI. Because REFUTES and REVISIONS represent different decision boundaries, the earlier review was not used as direct validation of the current benchmark labels. We instead report it as a characterization of expert agreement on the predecessor task. We conducted this analysis using the open-source text annotation tool doccano^65^. Pairs for review were selected by stratified random sampling in order to ensure pairs from each class label and partition (where applicable) were included in the chosen subset. We also aimed to maximize diversity in the source publication of the evidence, avoiding additional claims from the same publication (CIVIC may include multiple entries from the same publication if, for example, there are multiple mutations evaluated in the publication). We measured Inter-Annotator Agreement (IAA), the extent to which different annotators agree when labeling the same data, with Fleiss’ kappa score since we have more than 2 annotators. A subset of claim/evidence pairs were annotated with a written explanation of the stance label.

## Supplementary Results

### Historical expert-agreement results

Historical expert-review analyses were conducted using the predecessor SUPPORTS/REFUTES/NEI formulation and are reported separately because they do not directly validate the current REVISIONS category.

On the predecessor SUPPORTS/REFUTES/NEI task, expert annotators achieved a Fleiss’ κ^68^ of 0.65 and 77.9% agreement (p < 0.001). These results indicate substantial^69^ agreement for the earlier evidence-stance formulation but should not be interpreted as direct validation of the current REVISIONS category or all reference labels in the final release. Most disagreements involved NEI versus another label, reflecting differing thresholds for determining when a claim element could be inferred from the source text. The high agreement levels suggest that despite the complexity of biomedical literature, human experts can often reach consensus on claim veracity given appropriate context, providing a valuable benchmark for model performance. In nearly all cases (88.1%, Supplementary Fig 4a), disagreements between annotators were between NEI and one of the other labels. For example, annotators disagreed about when details of the claim could be reasonably inferred from the evidence or required more explicit mention in the source text (Supplementary Fig 4b).

### Prompt Optimization Improves Model Performance

Claim verification is defined as a model’s ability to correctly label the stance of evidence toward a claim when given both the claim and evidence as input. In order to get a measure of language models reasoning performance when the expected evidence was provided, we evaluated language models on the CIViC-Fact dataset with the manually selected evidence collected from hypothes.is or simulated examples (see methods).

It is computationally expensive to train or fine-tune a large language model, therefore recent work has explored ways to improve the output from a language model without fine-tuning or otherwise changing the actual model. Instead, we adjust the input to the model, this is called in-context learning (ICL). We tested a large range of open LLMs for claim verification, smaller models (Qwen 3.5 9B/27B; BioMistral; Llama 3.1 8B; Llama 3.2 1B/3B) performed poorly in initial experiments. Therefore, only Llama 3.2 [3B] was tested downstream as a comparator for the fine-tuned version of this model. A full list of models explored can be found in Supplementary Table 1. The 4 largest models, along with GPT 5.2, were chosen for further prompt optimization: Llama 3.1; Llama 4-Scout; Gemma 4; and Mixtral (Supplementary Fig 5). Both Llama 3.1 and Mixtral showed some improvement with the chat agent system prompt (Supplementary Fig 5a). Llama 3.1 and Llama 3.2 were the only models to show non-negligible improvement with the inclusion of the generated explanations (Supplementary Fig 5c, +7%). Nearly all models benefitted from additional examples (Supplementary Fig 5d). The largest variation was seen in changing the user prompt (Supplementary Fig 5b, Supplementary Table 1).

### Historical explanation-quality results

Given that the strength of using an LLM for classification lies in the reasoning capabilities, we separately evaluated explanations generated under the predecessor SUPPORTS/REFUTES/NEI task. Human experts tagged each response for helpful content and for factual, logical and communication-related problems (Table 1). Because this evaluation was conducted before the introduction of the REVISIONS category, the results assess the grounding and reliability of explanations for the original stance-classification task; they do not measure whether models can adequately explain or propose revisions under the current benchmark formulation. Some responses were particularly impressive. For example, Gemini 2.5 demonstrated strong logical reasoning by identifying a mistake in the variant name reported in the source publication. The claim listed the variant as S80N, while the publication included S80I, but in reality, all other representations in the source consistently referred to S80N. Gemini detected this discrepancy by translating the three-letter codon into its corresponding amino acid (Supplementary Fig 6a). At the same time, models also produced coherent but problematic answers. GPT-4.1, for instance, incorrectly claimed that ibuprofen was tested in the source publication; while the claim mentioned ibuprofen, it was not actually included among the therapies studied. Both Gemini 2.5 and GPT 4.1 also made logically plausible but ultimately incorrect inferences (Supplementary Fig 6b).

The overall inter-annotator agreement among tags was fair (FK: 0.30, PA: 60.4%, p < 0.001). When we calculate agreement on the union of the error tags the agreement is considerably higher (FK: 0.44, PA: 74.8%, p < 0.001), indicating that while there is some disagreement amongst annotators on the individual type of problem, annotators reach a greater consensus that a problem exists. The lowest agreement was found among untagged content (FK: 0.15, PA: 75.4%, p < 0.001) indicating greater disagreement amongst annotators on whether or not content was relevant. Given this level of agreement, each response, along with its initial tags, was re-examined by an additional annotator or collaboratively discussed by the group to resolve disagreements, and to capture additional tags such as “*incomplete”* and “*misleading, or overstated”*. All subsequent results are reported from this re-examined set of annotations. Most models produced responses with some helpful content, but the rates of problematic entries were also quite high (Supplementary Fig 7a).

GPT 4.1 performed the best in providing helpful responses but still had major issues (factual errors) in 14% of responses. Interestingly, despite being a completely different dataset, this rate of hallucinations is similar to previous work which observed a hallucination rate of 15% on the biomedicine subset of the HaluEval2.0 dataset with ChatGPT^70^. Despite this, proprietary models still considerably outperformed the smaller locally run Llama model. 84% of GPT 4.1 responses were considered helpful with no or only minor issues, compared to 39% of Llama responses (Supplementary Fig 7b).

### Supplementary Figures

**Supplementary Figure 1.**
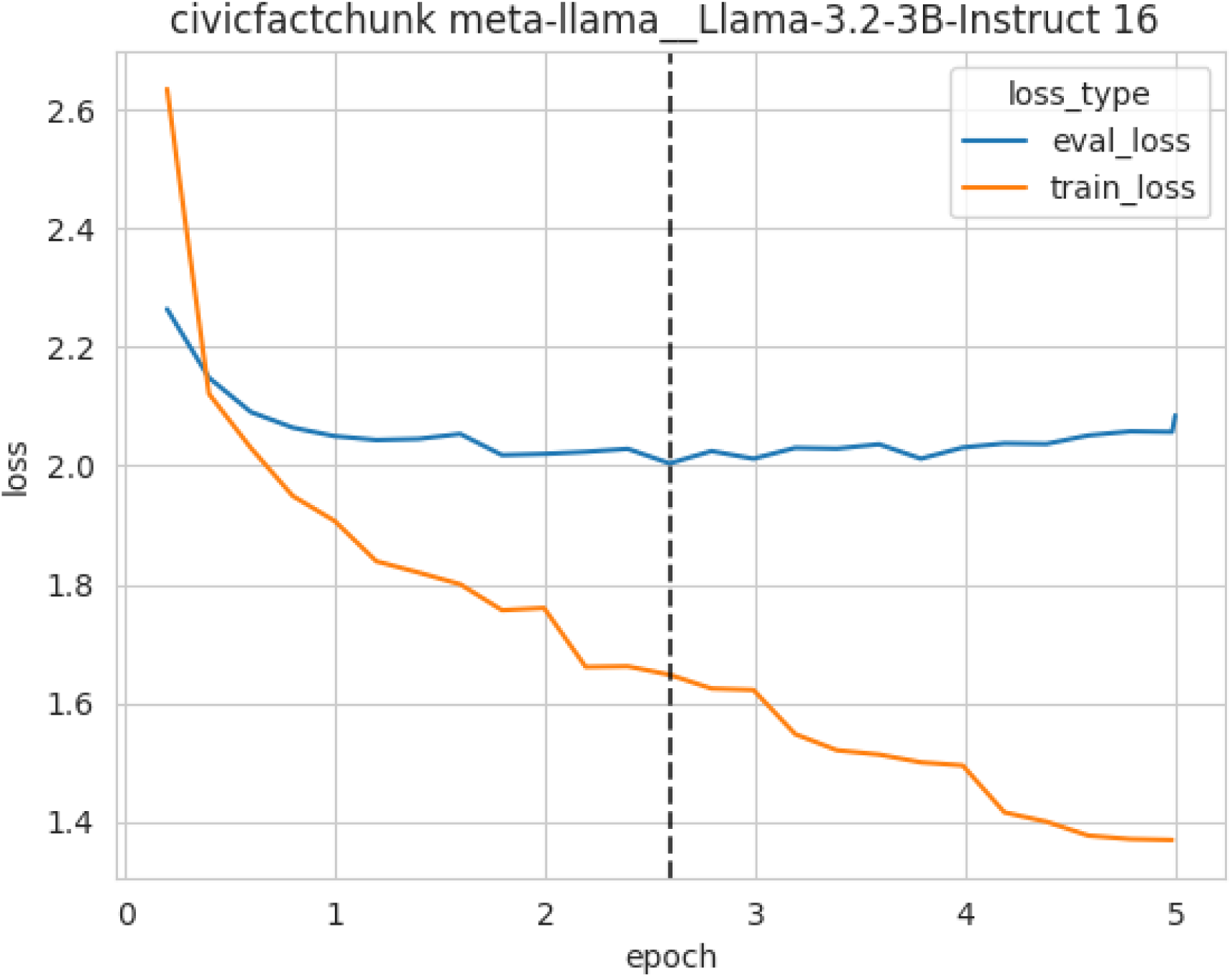
Training loss curve for LLama3.2 3B on the CIViC-Fact claim verification dataset. The model chosen by early stopping is marked with the dashed vertical black line.

**Supplementary Figure 2.**
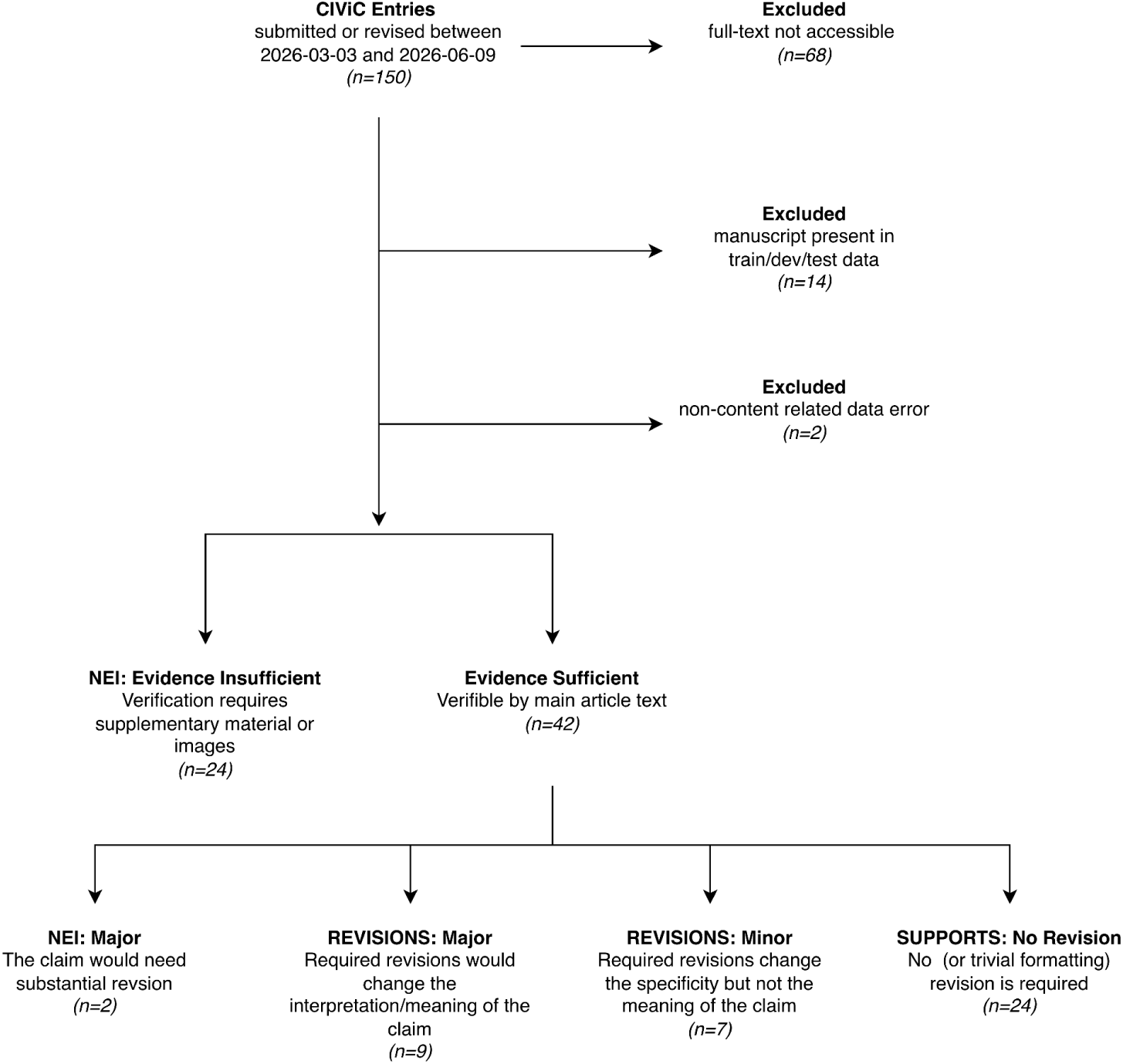
Selection of CIViC entries for the post-cutoff cohort.

**Supplementary Figure 3.**
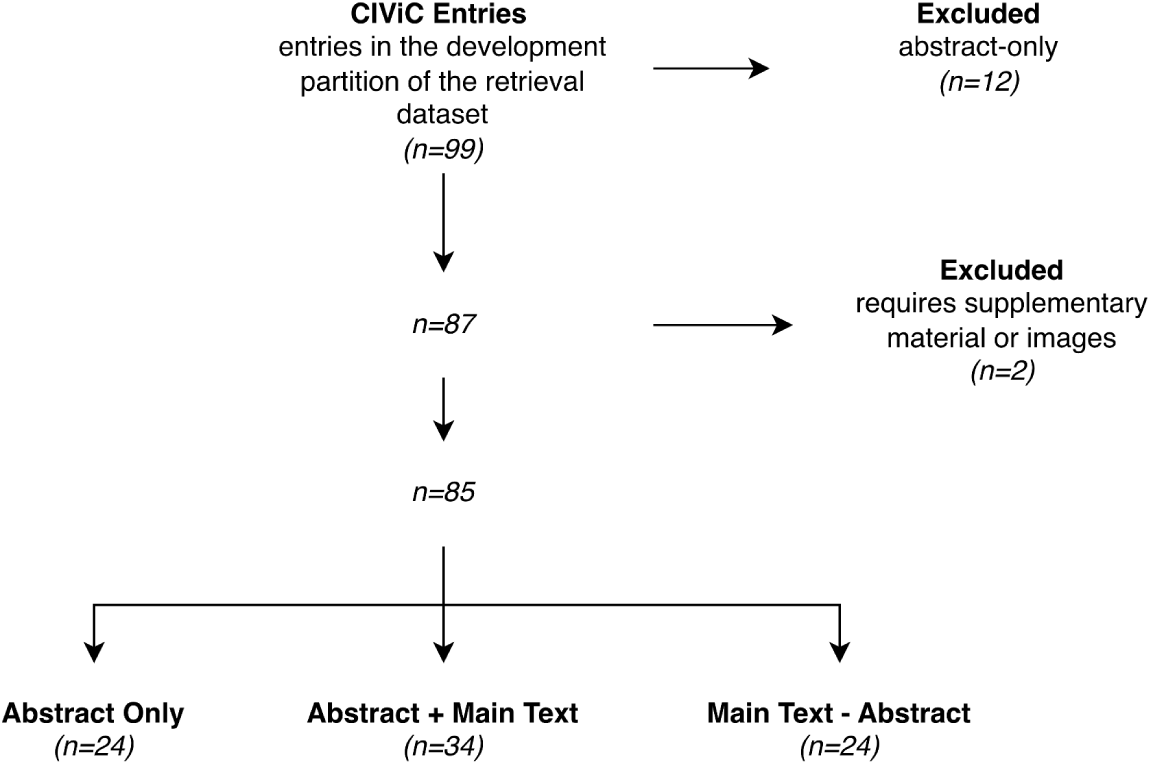
Distribution of evidence within the development partition of the retrieval dataset.

**Supplementary Figure 4.**
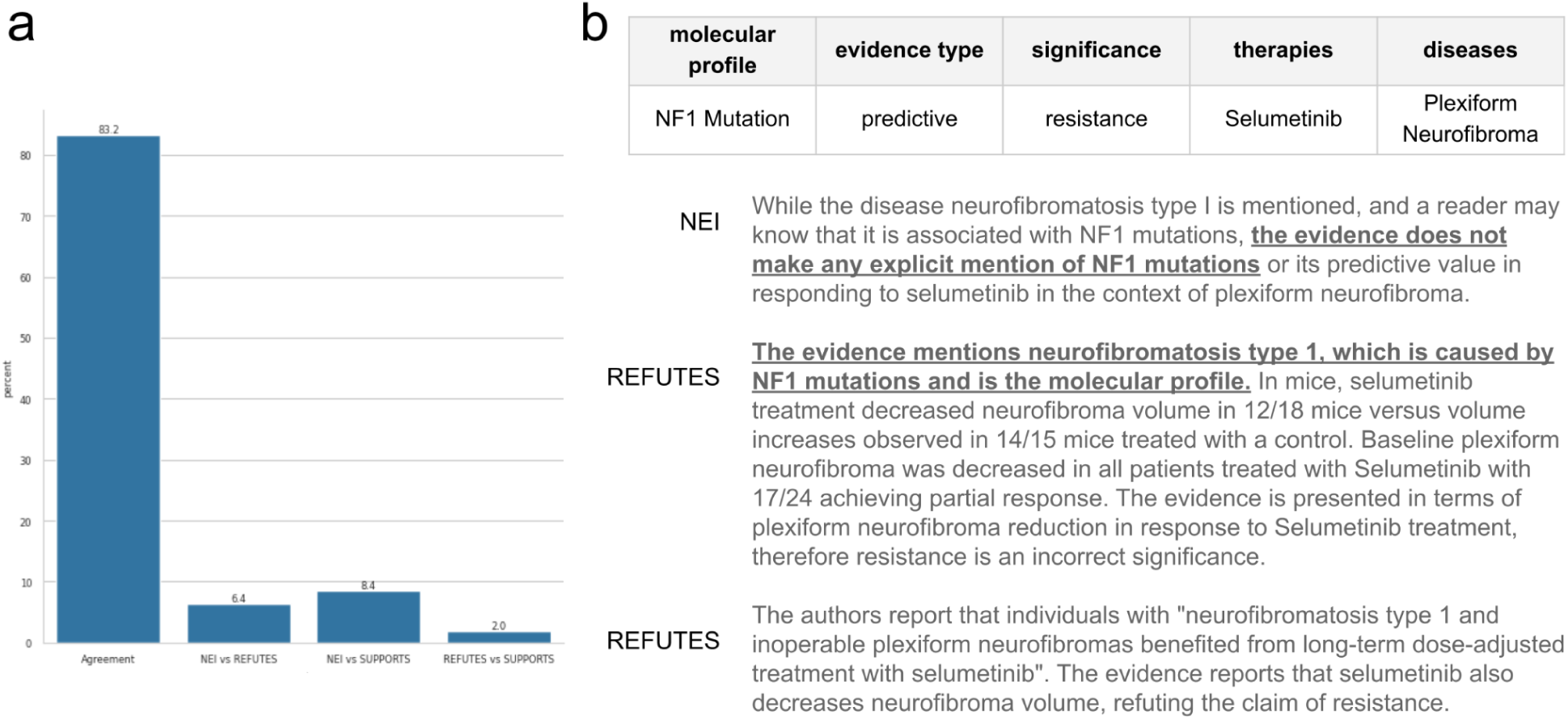
a. Proportion of total pairs of manually curated claim/evidence pairs with a given label. Disagreements are given as / . b. An example of disagreement among expert human annotators. The claim is given in the table at the top, evidence is not shown. Example of the stance label and explanation from three different annotators. The consensus label for this example is REFUTES. One annotator expects more explicit mentions in the source text while the other annotators agree it is sufficient to infer the presence of NF1 mutations from the disease. The evidence for this claim is omitted from the figure due to length. Minor typos have been corrected here for display, the raw reviewer commentary is available in the dataset.

**Supplementary Figure 5.**
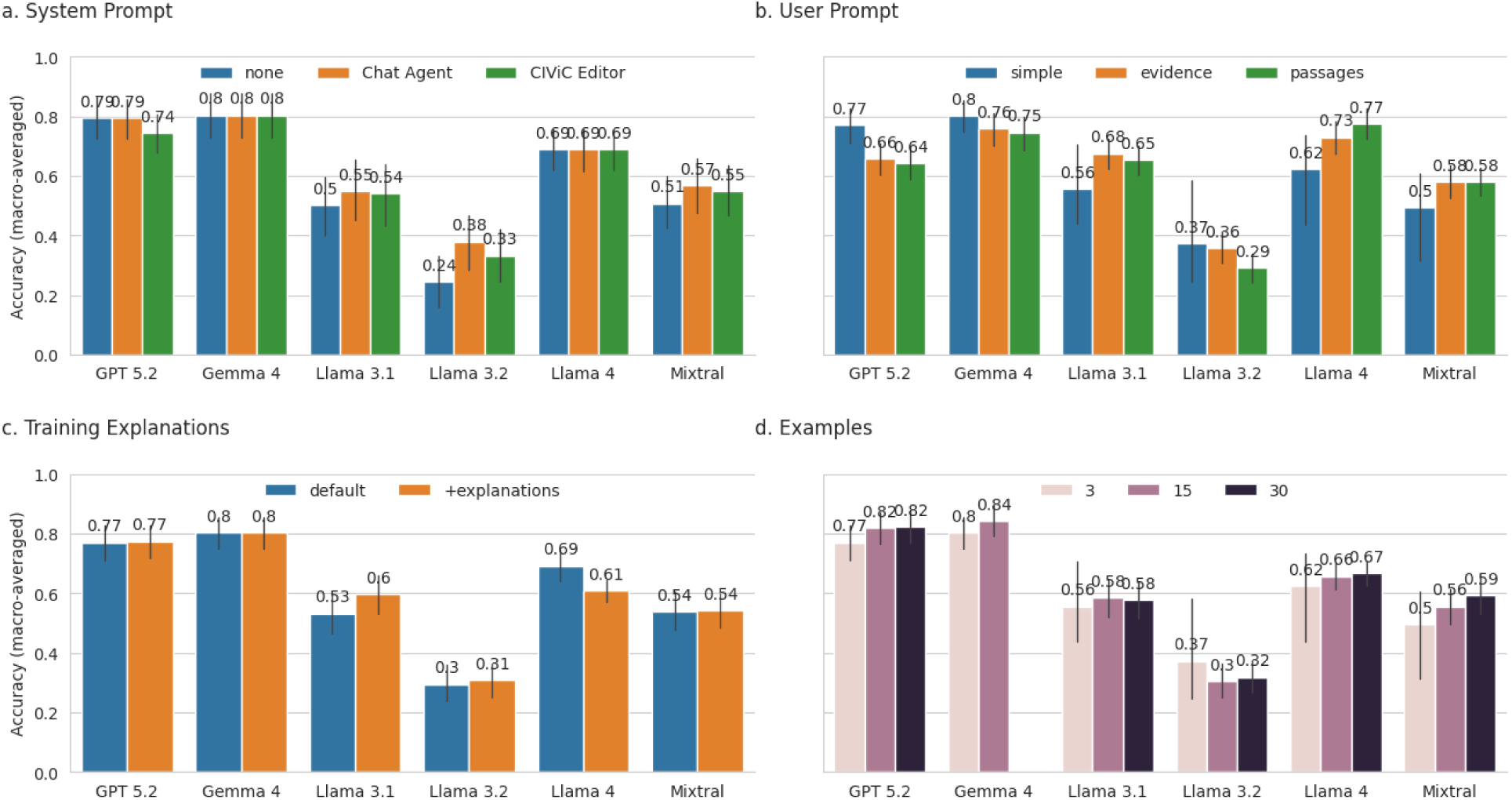
Prompt Optimization of few shot models. a. Model accuracy with different system prompts. Models were tested either with no system prompt or one of two system prompts. The first was generated by GPT5.5 pro to resemble a Chat agent in a similar style to what we expect from ChatGPT whereas the second “CIViC Editor” prompt was generated by reviewing the CIViC editor documentation. Both prompts are given in the supplementary material. b. The user prompt tested three different but related sets of user instructions where evidence was given simply as text (simple), or with more explicit institution and xml separation tags (evidence, passages). The evidence user prompt gave the evidence as a single block whereas the passages prompt distinguishes the individual passages and their position within the original document. c. Explanations were generated for the training data in the CIViC-Fact dataset using GPT5.2. We then tested whether including these explanations in the examples provided to the model affected accuracy. d. Models were tested with 3, 15, or 30 examples.

**Supplementary Figure 6.**
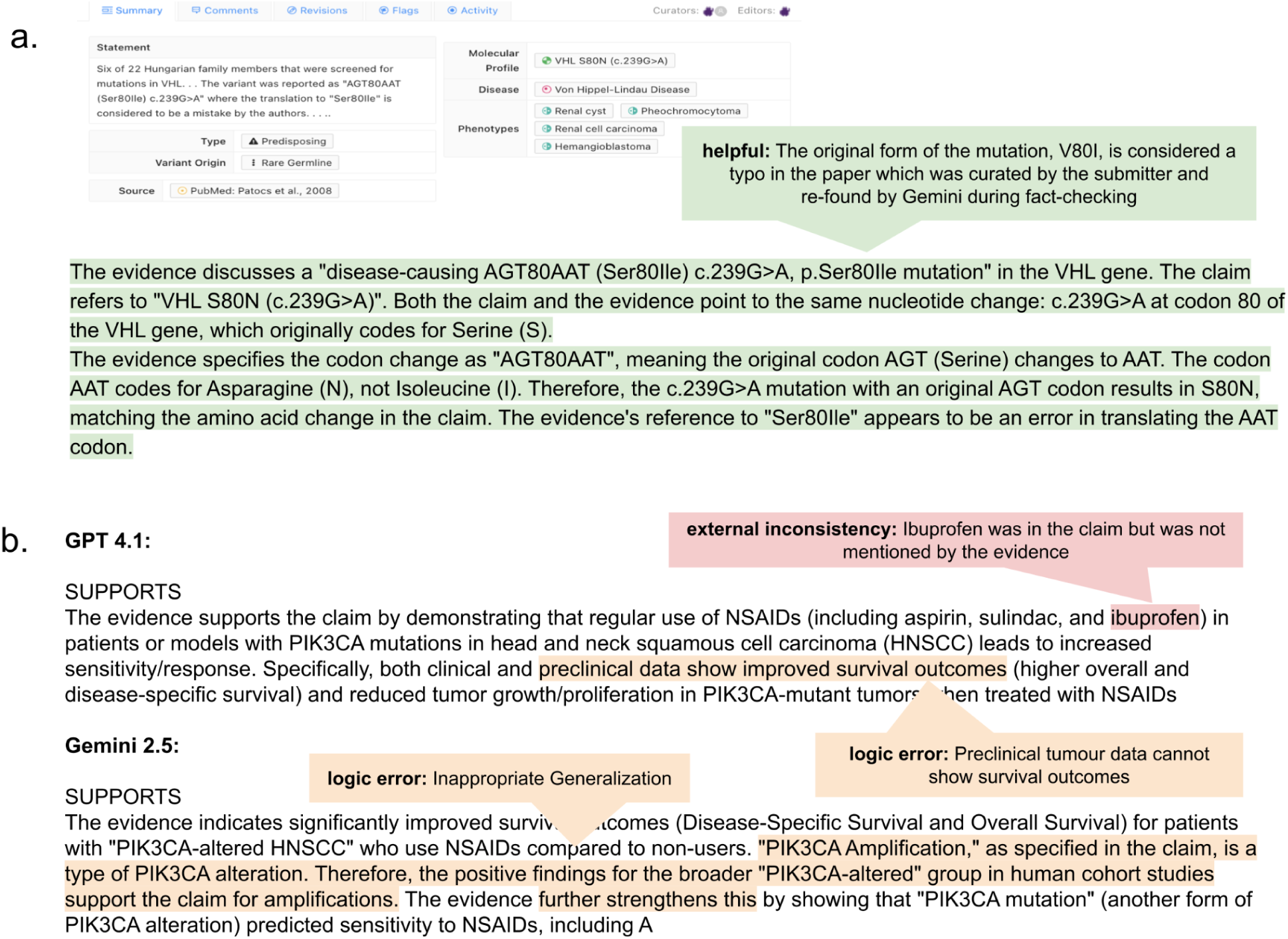
a. Gemini identified a typo in the original paper and used the three-letter codon and CDS notation to verify the variant matched the claim. b. Both Gemini and GPT produced major issues manually annotated by expert human annotators.

**Supplementary Figure 7.**
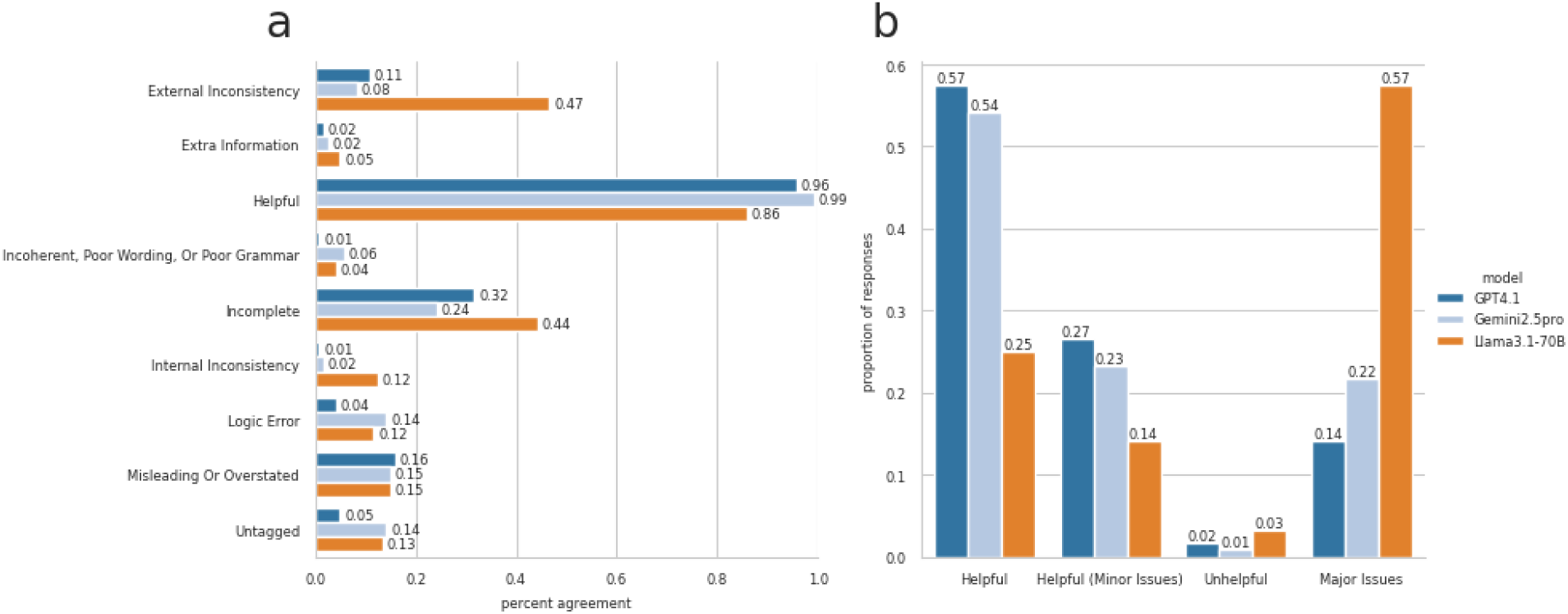
a. Individual rates of tags used by response author (generative model). b. Classification of responses. We categorized responses into four groups based on their tagging results: Major Issues, if any major issues were identified; Helpful (Minor Issues), if the response was considered helpful and only minor issues were tagged; Helpful, if the response was helpful with no issues identified; and Unhelpful for all remaining responses.

### Supplementary Tables

**Supplementary Table 1.** Prompting configurations chosen for Downstream Experiments. Note that due to high GPU VRAM usage Gemma 4 was only tested with 3 or 15 examples. All tests with 30 examples resulted in CUDA out of memory errors despite significant resources.

| Model | User Prompt | System Prompt | ICL Examples | Explanations |
| --- | --- | --- | --- | --- |
| Llama 3.1 70B | evidence (+12%) | Chat Agent (+5%) | 15 (+2%) | yes (+7%) |
| Llama 4 Scout | passages (+15%) |  | 30 (+5%) |  |
| Mixtral | evidence (+8%) | Chat Agent (+6%) | 30 (+4%) |  |
| GPT 5.2 | simple |  | 30 (+5%) |  |
| Gemma 4 | simple |  | 15 (+4%) ** |  |
| Llama 3.2 [3B] | simple | Chat Agent (+5%) | 3 | yes (+1%) |

